# Structural basis of *S*-adenosylmethionine-dependent Allosteric Transition from Active to Inactive States in Methylenetetrahydrofolate Reductase

**DOI:** 10.1101/2023.12.14.571224

**Authors:** Kazuhiro Yamada, Johnny Mendoza, Markos Koutmos

## Abstract

Methylenetetrahydrofolate reductase (MTHFR) is a pivotal flavoprotein connecting the folate and methionine methyl cycles, catalyzing the conversion of methylenetetrahydrofolate to methyltetrahydrofolate. Human MTHFR (*h*MTHFR) undergoes elaborate allosteric regulation involving protein phosphorylation and AdoMet-dependent inhibition, though other factors such as subunit orientation and FAD status remain understudied due to the lack of a functional structural model. Here, we report crystal structures of *Chaetomium thermophilum* MTHFR (*c*MTHFR) in both active (R) and inhibited (T) states. We reveal FAD occlusion by Tyr361 in the T-state, which prevents substrate interaction. Remarkably, the inhibited form of *c*MTHFR accommodates two AdoMet molecules per subunit. In addition, we conducted a detailed investigation of the phosphorylation sites in *h*MTHFR, three of which are novel. Based on the structural framework provided by our *c*MTHFR model, we propose a possible mechanism to explain the allosteric structural transition of MTHFR, including the impact of phosphorylation on AdoMet-dependent inhibition.

## Introduction

Methylenetetrahydrofolate reductase (MTHFR) catalyzes the enzymatic reduction of methylenetetrahydrofolate (CH_2_-H_4_folate) using NAD(P)H as the reducing agent^1,2^. MTHFR serves as the major source of methyltetrahydrofolate, which is the substrate of the downstream enzyme, methionine synthase^3^. Methionine synthase catalyzes the formal methyl transfer from methyltetrahydrofolate to homocysteine, yielding tetrahydrofolate and methionine as products^4,5^. Methionine is thereafter converted to *S*-adenosylmethionine (AdoMet), which is known as biology’s universal methyl donor. In addition, AdoMet acts as an allosteric regulator of MTHFR and feedback loop regulator in folate and methionine metabolism (Fig. 1a)^6, 7, 8^. The regulation of MTHFR activity via AdoMet is crucial in controlling the flux of folate-carrying one-carbon units to methionine synthesis, helping bridge the folate and methionine cycles, as it is the only known enzyme to exhibit AdoMet-dependent inhibition^9^.

**Figure 1.**
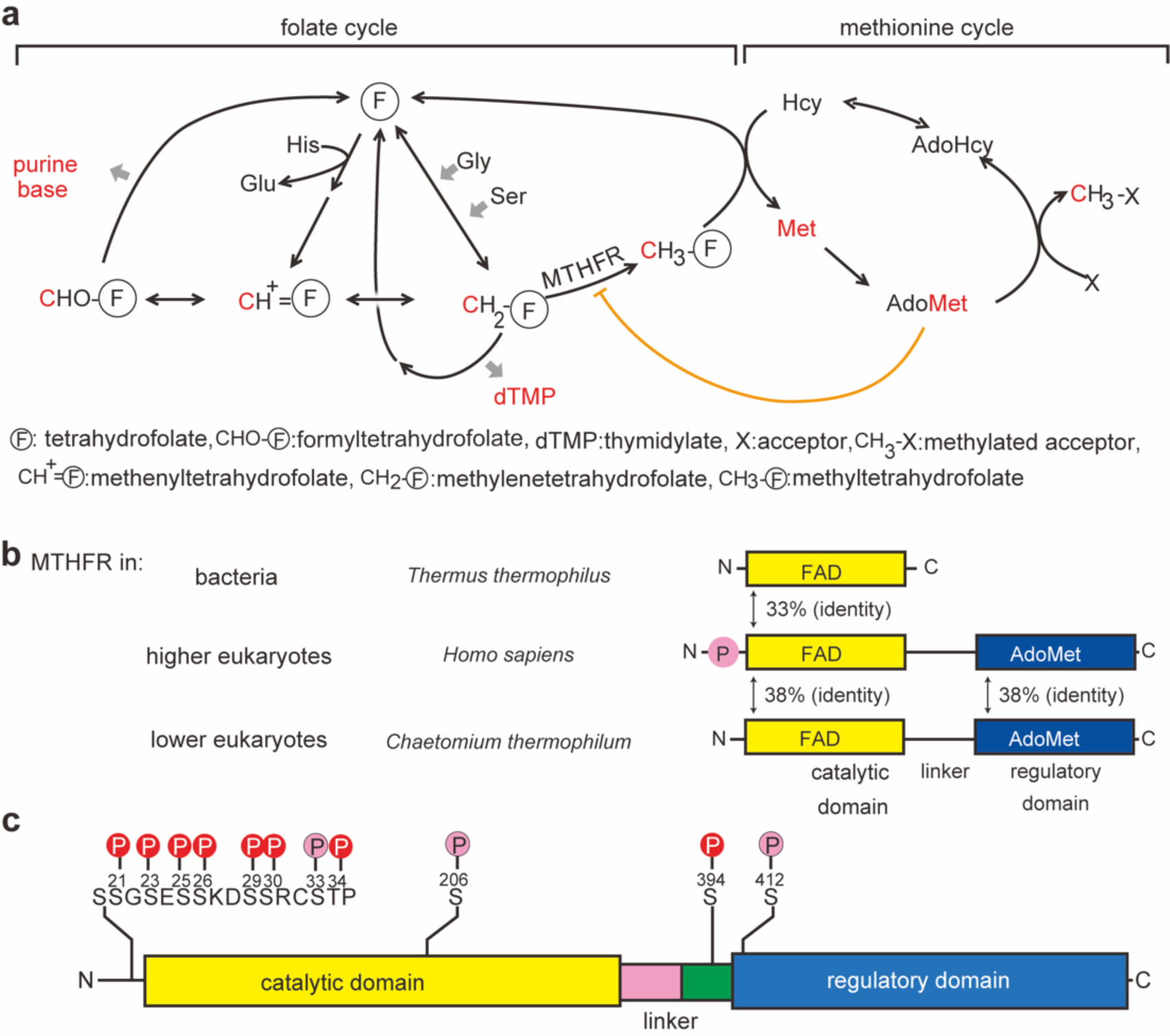
Folate and methionine metabolism one-carbon cycle. **a** C-units (such as formyl, methenyl, methylene, or methyl group) carried by folate are supplied from amino acids (Gly, Ser, and His). C1-units and metabolites incorporating these one-carbon units are shown in red. C1-units are used to synthesize purine bases, dTMP, and methionine (Met). Methionine is synthesized to AdoMet, which is the major methyl donor in the cell. AdoMet is an allosteric inhibitor for MTHFR, whose activity determines the flux of folate C1-unit to methionine synthesis and transmethylation. **b** Domain organization of MTHFRs. The catalytic domain of MTHFR contains FAD as an essential cofactor (shown in yellow). Although bacterial MTHFRs only consist of catalytic subunits, eukaryotic MTHFRs consist of both a catalytic domain and a regulatory domain, connected by a linker. AdoMet binds to the regulatory domain. Human (*Homo sapiens*) MTHFR has a Ser/Thr-rich region on its N-terminus which can be phosphorylated (shown in circled-P with pink), enhancing its sensitivity for allosteric regulation by AdoMet. In this study, MTHFR from *Chaetomium thermophilum* is used as a structural model for *h*MTHFR, with which it shares reasonable homology. **c** Phosphorylation mapping of *h*MTHFR. Phosphorylation sites were identified by our analysis using tryptic fragments and LS-MS/MS analysis (indicated by circled P in pink). Previously reported sites are indicated by the red lines. Overlapping phosphorylation sites in both results, which occurred on eight amino acid side chains, are represented by red circled P. Phosphorylation occurs intensively in the N-terminal Ser-Thr rich site and the C-terminal end of the linker.

MTHFR non-covalently binds FAD as an essential cofactor, shuttling electrons from NAD(P)H in a ping-pong bi-bi mechanism^10^. The eukaryotic form of MTHFR is homodimeric, where each monomer consists of the N-terminal catalytic domain connected by a linker to the C-terminal regulatory domain (Fig. 1b)^7^. In contrast, bacterial MTHFR lacks the regulatory domain and manifests as a homooligomer consisting only of catalytic subunits^11,12,13^. The catalytic component of all MTHFRs bind FAD within the core of a β_8_α_8_ barrel structure, also known as a TIM-barrel. The regulatory module has a unique structural topology that facilitates the formation of MTHFR dimers through the interface of its regulatory domains^7^. This domain has been shown to bind *S*-Adenosyl-L-homocysteine (AdoHcy) and is presumed to bind AdoMet, though no structure of the AdoMet bound form currently exists.

MTHFR has evolved several forms of (allosteric) regulation, ranging from substrate-induced inhibition in bacteria (NADH)^12^, NADH preference in bacteria as opposed to NADPH preference in eukaryotes^14,15,16^, to the more complex AdoMet-dependent allosteric inhibition observed in eukaryotes^7^. This AdoMet-dependent allosteric regulation is unique to eukaryotic MTHFRs, as the regulatory domain thought to be responsible for AdoMet-binding/sensing is not present in bacteria. However, to date, no structure of the inactive enzyme (T-state) exists, despite a relative wealth of biochemical data supporting an important role of AdoMet in regulating MTHFR activity^6,7,8,9,14,17^. Pioneering studies conducted in 1987 highlight that the eukaryotic MTHFR conformational ensemble exists in an R vs. T-state equilibrium, even as an apoenzyme (K_RT_, ([T]/[R] in absence of ligands = 0.30)^17^. AdoMet acts as an allosteric inhibitor, orchestrating and facilitating the structural transition to the inhibited conformation (T-state), while NADPH assumes the role of an activator, shifting the conformational equilibrium to the activated conformation (R-state)^17^. While AdoHcy does not act as an activator per se, it does act as a competitive binder for AdoMet^1^. However, the lack of a structural blueprint of eukaryotic MTHFR, particularly in its inactive, T-state configuration, prevents interrogating the underlying molecular mechanism that dictates the dynamics of the allosteric transition.

In this report, we have used a thermophilic MTHFR homolog from *Chaetomium thermophilum* (*c*MTHFR) as a structural and biochemical model for *h*MTHFR. Although *c*MTHFR lacks the Ser-rich N-terminal phosphorylation region, the catalytic and regulatory domains of *c*MTHFR share high levels of sequence conservation/homology with those of *h*MTHFR (Fig. 1b, 38% sequence identity), allowing us to investigate allosteric regulation *independent* of phosphorylation status. Biochemical analysis of *c*MTHFR confirmed its NADPH-preference and AdoMet-mediated inhibition, two features common to eukaryotic MTHFRs. Structural elucidation and recapitulation of the previously solved active, R-state conformation^7^ displays the validity of our biochemical and structural model. Additionally, by introducing a human patient mutation, Arg357Cys (Arg315 in *c*MTHFR), we successfully crystallized *c*MTHFR in its inhibited, T-state, AdoMet-bound confirmation. To complement and further understand the allosteric transition between each state, a detailed investigation of the phosphorylation sites in *h*MTHFR was conducted. Here, we discuss the possible shared mechanism of the allosteric transition of eukaryotic MTHFRs.

## Results

### Biochemical Characterization of *h*MTHFR – Phosphorylation Site & Patient Mutation Analysis

In eukaryotes, MTHFR is a homodimer consisting of catalytic and regulatory domains. Notably, the only structure of a eukaryotic MTHFR (*h*MTHFR, 6FCX) shows that the dimer interface consists of interactions between the regulatory domains of two different protomers that form an antiparallel β-sheet dimer/β-sandwich. In this structure, AdoHcy was bound to the regulatory domain, and FAD was bound in the catalytic active site in a solvent-exposed manner, indicating that this captured *h*MTHFR structure likely represented a catalytically-competent, active enzyme in the R-state.

Human (*Homo sapiens*) MTHFR (*h*MTHFR) is post-translationally modified via phosphorylation of several residues, most of which reside on its N-terminus and the bulk of which are Ser. Phosphorylation of most sites is dependent on the initial phosphorylation of Thr34 in *h*MTHFR, which is thought to be the priming position for post-translational modification of *h*MTHFR by Pro-directed kinase(s)^18^. Phosphorylation plays a role in AdoMet-dependent allosteric regulation of MTHFR as indicated by experiments showing that mutation of the homologous Thr residue to Ala in other eukaryotic MTHFR proteins renders the enzyme less sensitive to AdoMet binding and inhibition^14,17^. However, the structural details of how the N-terminal phosphorylation sites affect the R-vs T-states conformational ensemble and how AdoMet binding leads to allosteric inhibition of *h*MTHFR are not known.

LC-MS/MS analysis of *h*MTHFR showed that, in our *h*MTHFR preparations, eleven amino acids were phosphorylated (Ser21, Ser23, Ser25, Ser26, Ser29, Ser30, Ser33, Thr34, Ser206, Ser394, and Ser412), eight of which overlapped with a previous report and three of which are new (Ser33, Ser206, Ser412). The previous report found 16 total amino acids were phosphorylated, 11 of which were found on the N-terminus^7^. The identification of Ser33 as one of the phosphorylation sites in the N-terminal Ser/Thr rich region, not previously reported, is noteworthy. Previously, Ser33 was postulated to play a role in phosphorylation based on data that showed that a Ser33Ala mutation impedes phosphorylation of other confirmed phosphorylation sites in *h*MTHFR^14^, suggesting the importance of Ser33 phosphorylation in the early stages of sequential phosphorylations of *h*MTHFR. Our analysis reveals three phosphorylation sites outside the N-terminal Ser/Thr rich region (Fig. 1c, Supplementary Figs. 1 and 2). While phosphorylation of Ser394 was previously established, Ser206 and Ser412 emerge as novel findings. The functional implications of phosphorylation at Ser206, Ser394, and Ser412 and their role in modulating *h*MTHFR activity remains unexplored.

The N-terminal sites were shown to be proximal to the linker in the R-state^7^. As alluded to previously, phosphorylation favors T-state formation by sensitizing AdoMet allosteric inhibition^14^, and conversely, dephosphorylation desensitizes AdoMet-mediated allosteric inhibition. The role of the linker remains unclear, especially in the R- to T-state transition, as it has only been observed in the R-state. The linker is thought to interact with the phosphorylated amino acid side chains of the N-terminal, albeit in a differential mode depending on the conformation (R vs. T-state).

Despite the discovery of three new phosphorylation sites in addition to confirming the phosphorylation of eight previously identified residues, *h*MTHFR was difficult to study structurally and recalcitrant to crystallization in our hands. Encouraged by the recapitulation of the results of previous phosphorylation studies, and taking a cue from patient mutations in *h*MTHFR^19^, we set out to see if we could rationally trap the elusive T-state. Using the NADPH-menadione oxidoreductase assay, the effects of several patient mutations on activity and FAD release were analyzed and compared to wild-type *h*MTHFR (Supplementary Fig. 3) to discern any trends that would provide insights into the catalytic mechanism or the conformational rearrangements influenced by AdoMet binding.

Of all the patient mutations analyzed, Arg357Cys stood out due to its near abolishment of activity (14% relative to wildtype), *without* any alteration in FAD retention, as compared to the wild type *h*MTHFR (Supplementary Fig. 3). Previous studies in *h*MTHFR have shown that AdoMet binding leads to a corresponding *decrease* in FAD release^20^, indicating that the biochemical and, by extension, structural properties of the Arg357Cys mutant follow an analogous trend to those of the AdoMet-bound and inhibited enzyme in the T-state. In all, these results offer the tantalizing possibility that Arg357Cys favors the elusive inactive T-state form, mimicking the same conformation induced by AdoMet binding.

Thus far, despite numerous attempts, we have been unable to obtain structures of *h*MTHFR in its active or inactive conformations. However, given the promising biochemical data indicating that the T-state could be rationally trapped, we pursued alternate biochemical and structural models of *h*MTHFR among other homologs. Our intent was to elucidate the structural mechanism behind AdoMet-mediated allosteric inhibition while granting greater insights into the intricacies involved in the conversion of chemical signals into structural rearrangements. Consequently, we explored an MTHFR homolog from *Chaetomium thermophilum* (*c*MTHFR), a thermophilic fungus, as a structural and biochemical model for eukaryotic MTHFRs. Although *c*MTHFR lacks an N-terminal phosphorylation site, the catalytic and regulatory domains of *c*MTHFR share reasonable homology with those of *h*MTHFR (Fig. 1b, 38% sequence identity). Moreover, the lack of an N-terminal phosphorylation region in *c*MTHFR was a desired feature, selected to facilitate crystallization through increased homogeneity, uncoupling phosphorylation status from effector-induced allosteric inhibition and structural transitions.

### *c*MTHFR as a Biochemical Model for Eukaryotic MTHFR

We first set out to validate *c*MTHR as a functional biochemical model for *h*MTHFR and eukaryotic MTHFRs in general. We determined its reductant preference using the NAD(P)H-menadione oxidoreductase assay (Fig. 2a). *c*MTHFR^wt^ can oxidize both NADPH and NADH *but* prefers NADPH (120 µM/min) over NADH (44 µM/min), akin to previous studies of eukaryotic MTHFR models^21^. Having confirmed the NADPH-preference of *c*MTHFR^wt^, we set out to determine the effect of AdoMet on *c*MTHFR activity, again using the NAD(P)H-menadione oxidoreductase assay (Fig. 2a). In the presence of AdoMet (100 µM), activity was inhibited by 25%. The Arg357Cys patient mutation was translated into *c*MTHFR^wt^, corresponding to the *c*MTHFR^R315C^ mutant, and its NAD(P)H-preference and AdoMet-mediated inhibition were likewise interrogated. At the same enzyme concentration and without AdoMet, the Arg315Cys mutant consumed NADPH at a much slower rate than the wild-type enzyme (14% relative activity) (Fig. 2b), indicating that it existed in a primarily inhibited state.

**Figure 2.**
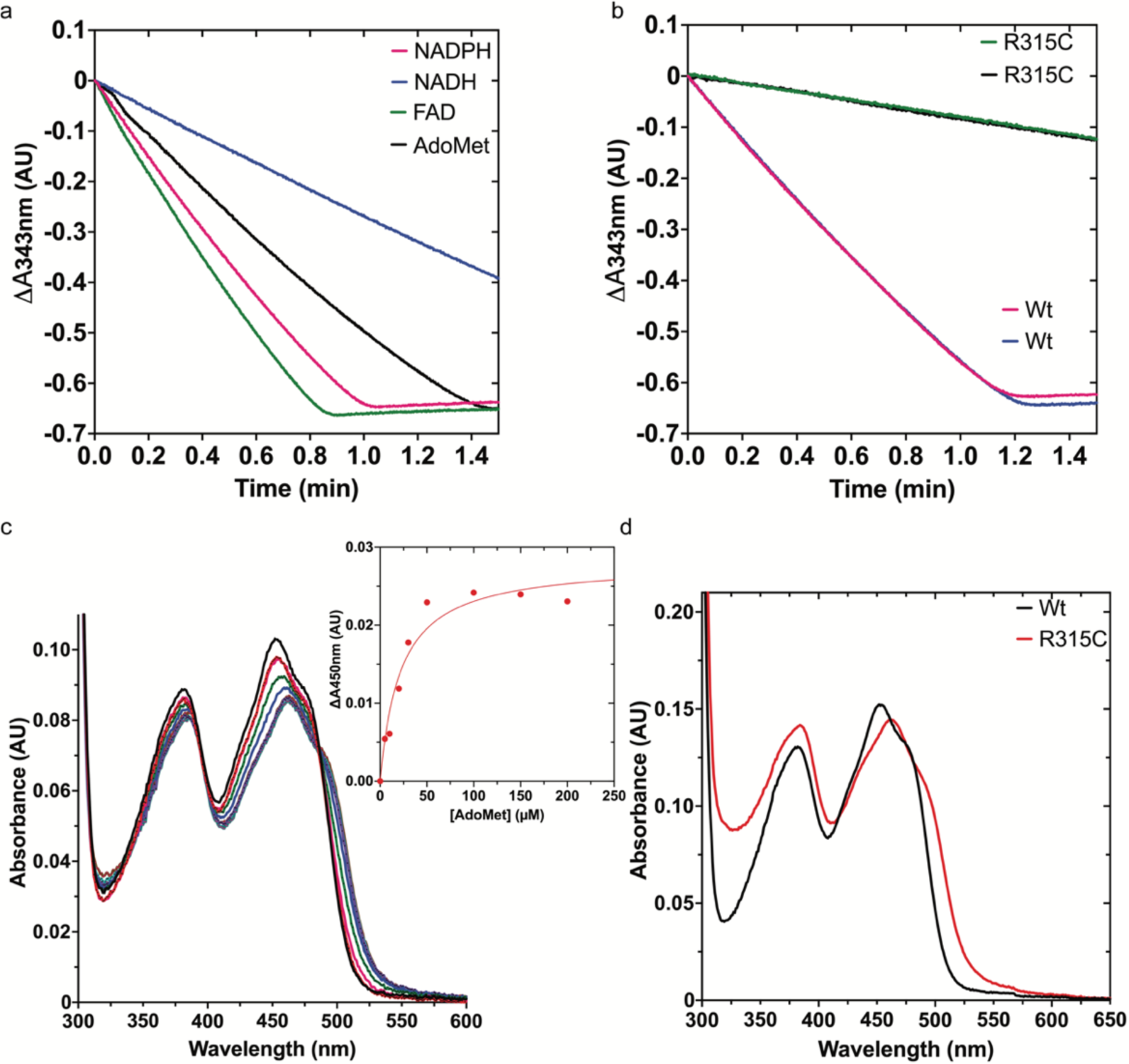
Initial biochemical characterization of *c*MTHFR. **a** NADPH/NADH preference and AdoMet-dependent inhibition of *c*MTHFR was assessed using the NADPH:menadione oxidoreductase assay. Consumption of NADPH or NADH was monitored by absorbance at 343 nm at room temperature. **b** NADPH-menadione oxidoreductase *c*MTHFR^R315C^ activity assayed in the same conditions as in panel **a**. **c** UV-Vis spectral changes of *c*MTHFR^wt^ titrated with AdoMet. The typical flavin spectrum of *c*MTHFR without AdoMet has two peaks at 370 and 450 nm. When AdoMet is added, these peaks are red-shifted and the absorbance at 450 nm is decreased. The inset shows the absorbance changes at 450 nm as a function of AdoMet. **d** UV-Vis spectra of *c*MTHFR^wt^ and *c*MTHFR^R315C^ in the absence of AdoMet. Source data are provided as a Source Data file.

The binding affinity of *c*MTHFR^wt^ titrated with AdoMet was monitored via spectral changes using UV-Vis (Fig. 2c), yielding a K_d,app_ of ∼23 µM. The UV-Vis spectrum of *c*MTHFR^wt^ is noticeably altered upon AdoMet binding, with FAD undergoing an AdoMet concentration-dependent absorbance quench and red-shift at 450 nm, altering the absorbance maxima of FAD from 453 nm to 463 nm. These absorbance changes are likely due to changes in the FAD microenvironment upon AdoMet binding (Fig. 2c). Spectra of *c*MTHFR^wt^ and *c*MTHFR^R315C^ in the absence of AdoMet were compared (Fig. 2d). The spectrum of *c*MTHFR^R315C^ is similar to that of *c*MTHFR^wt^ *with* AdoMet bound (Fig. 2d) (maxima 462 nm); therefore, we posit that the Arg315Cys mutation results in a “locked” MTHFR conformation that is similar to that induced by the AdoMet-binding, indicating that the *c*MTHFR^R315C^ mutant results in a conformation that resembles the elusive T-state. Encouraged by the biochemical results for both the *c*MTHFR^wt^ and *c*MTHFR^R315C^ mutant, we decided to test the validity of *c*MTHFR as a structural model, initially attempting to capture and recapitulate the R-state structure observed for *h*MTHFR.

### *c*MTHFR as a Structural Model for Eukaryotic MTHFR – R-State

*c*MTHFR was crystallized and its structure was solved to 3.4 Å (Fig. 3a) in the active R-state. Electron density for FAD was observed in every catalytic active site. Although electron density consistent with AdoHcy was found in the regulatory domain, it was weak and could not be faithfully modeled. Even so, the homodimer assembly was found to display the same dimer interface mediated by the regulatory domain, a β-sheet sandwich consistent with the one previously observed for AdoHcy-bound *h*MTHFR (R-state)^7^ (Figs. 3a, b). Unlike bacterial MTHFR homologs, which use their catalytic domains for oligomeric assembly, *c*MTHFR, like *h*MTHFR, relies on its regulatory domain for oligomerization. The catalytic domains were also found to face “away” from the dimer interface and each other and are not interacting with one another, in stark contrast to the direct role they play in dimerization in bacterial MTHFRs. In this active R-state, the *si*-face of FAD is unoccluded in the active site and forms an extended hydrogen-bonding network between N5 of FAD centered on conserved residues His81 and Asp50 (His127 and Asp92 in *h*MTHFR), along with a strong hydrogen bond between the universally conserved Thr52 (Thr94 in *h*MTHFR) and O4 of FAD (Supplementary Fig. 4). Given that the active site is solvent exposed and that access to FAD is unoccluded, the captured structure likely represents a catalytically-competent form of the enzyme in the active R-state, much like that observed for *h*MTHFR. Indeed, PISA analysis shows that while ∼75.95% of the FAD active site binding interface is buried, a channel/tunnel exists, accessible to the bulk solvent, close to the adenine moiety and with direct access to the *si*-face of FAD.

**Figure 3.**
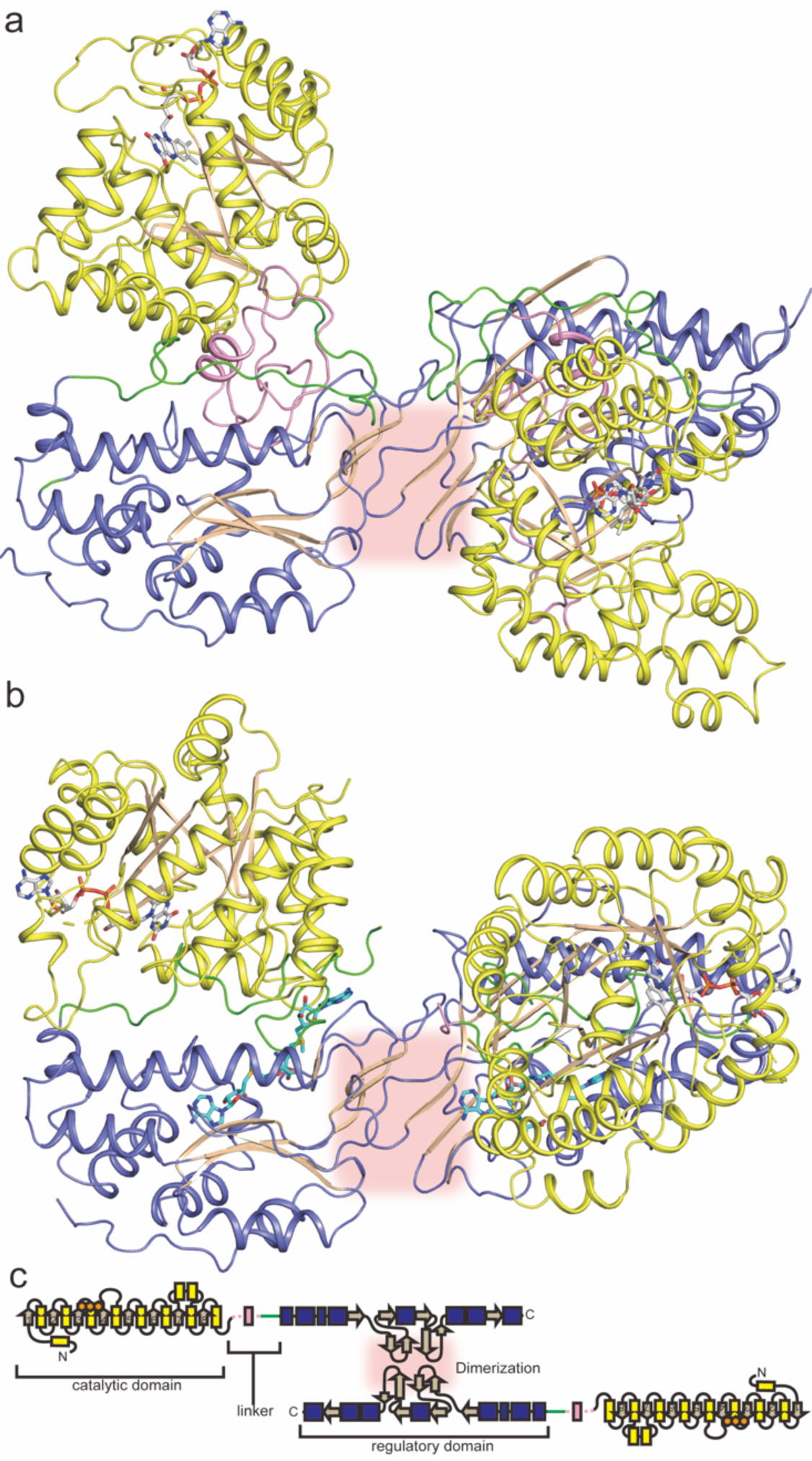
Dimer structure of *c*MTHFR. **a** Eukaryotic MTHFR structure in the R-state is illustrated in the ribbon diagram. The catalytic domain and regulatory domain are colored yellow and blue, respectively. The linker’s N-terminal region is shown in pink, while the C-terminal one is shown in green. The dimer interface is highlighted with a red box. **b** Eukaryotic MTHFR structure in the T-state is shown in the same orientation as that in the R-state (Panel **a**). **c** Schematic diagram of the secondary structural features in the dimeric eukaryotic MTHFR structures.

MTHFR conformational changes have been previously analyzed by limited tryptic digestion^8,14^. Matthews *et al.*^22^ previously demonstrated that porcine liver MTHFR (molecular weight of 77 kDa) undergoes cleavage into two major fragments of 39 kDa and 36 kDa, facilitated by limited proteolysis using trypsin. Conformational changes of *c*MTHFR, in the presence or absence of AdoMet, were visualized by limited proteolysis followed by SDS-PAGE (Supplementary Fig. 5). *c*MTHFR exhibits increased susceptibility to trypsin in the presence of AdoMet; this suggests that at least part of the linker, nestled between the catalytic and regulatory domains as evident in the R-state (Supplementary Fig. 5a), becomes solvent exposed in the T-state. The differences in the protease accessibility of the linker can be possibly attributed to an expected steric clash of Arg326 on the N-terminal region of the linker with the methyl group of AdoMet upon its binding (Ala368 in *h*MTHFR)^7^. This and possible local changes in the electrostatics could trigger linker rearrangement and transition to the T-state, where part of the linker becomes accessible. However, while the rich biochemical data available indicate that MTHFR exists as a conformational ensemble with unique allosteric properties that do not ascribe to typical models of allostery (such as the fast-lag phase of AdoMet inhibition and biphasic inhibition nature)^1^, the structural transition between the uninhibited (R-state) and AdoMet-bound inhibited (T-state) remains the subject of speculation given the lack of a structure in the T-state.

### Capturing the elusive T-state configuration using rational protein engineering

Encouraged by the recapitulation of the previously captured *h*MTHFR R-state structure, we set out to capture the elusive T-state structure. Taking a cue from our biochemical analysis of *h*MTHFR patient mutations, we set out to see if we could rationally trap the T-state using the Arg357Cys patient mutant (Arg315Cys in *c*MTHFR). As compared to the other patient mutants analyzed, the combination of FAD retention *and* catalytic activity abolishment (Supplementary Fig. 3), along with biochemical properties that mimic that of the AdoMet-bound state (Fig. 2), led us to focus on trapping the T-state via mimicry of the Arg357Cys patient mutant.

The analogous Arg315Ala mutant was used for structural studies to avoid the possible introduction of undesirable inter-disulfide bonds introduced by a Cys mutation; the *c*MTHFR^R315A^ mutant was successfully crystallized, with its structure solved to 2.8 Å (Fig. 3c). In this inactive T-state, the *si-*face of FAD is occluded by Tyr361 on the C-terminal region of the linker, which forms a π-stacking interaction (Supplementary Fig. 6). Additionally, each monomer adopts a more compact structure relative to the R-state, where the catalytic domain is rotated by 45° coupled to a translation of ∼16 Å, burying the catalytic domain between the linker and the regulatory domain. PISA analysis shows that ∼95.53% of the FAD active site binding interface is buried, aided by the new interaction from the linker. The regulatory domain was found to bind AdoMet, as expected for the T-state. Unexpectedly, two molecules of AdoMet were found per regulatory domain. One is found bound in the same site that AdoHcy occupies in the R-state (AdoMet site-1), while the other is found in a second adjacent site (AdoMet site-2) that was occluded by residues of the linker (aa301-305) in the R-state structure (Supplementary Figs. 7 and 8).

### Linker Rearrangement Dictates and Defines R- to T-State Transition

The overall structure of the *c*MTHFR^R315A^ mutant is shown in Fig. 3c. The dimer interface of *c*MTHFR^R315A^ in the T state is similar to that of the *h*MTHFR structure (6FCX) and *c*MTHFR in the R-state, mediated by the same β-sandwich formation between two monomers. Notably, the catalytic domains remain free of direct interdomain contact within the dimeric assembly. The superposition of *c*MTHFR^R315A^ in the inactive T-state with the *c*MTHFR structure in the active R-state using the regulatory domain as the reference point is shown in Fig. 4a. While the catalytic and regulatory domains of each state are indistinguishable (Supplementary Fig. 9), the global rearrangement between states highlights that the linker plays a focal role in channeling the R- to T-state transition. Indeed, the linker is found to be more flexible in the T-state and partially solvent-exposed, with part of the N-terminal region unable to be modeled.

**Figure 4.**
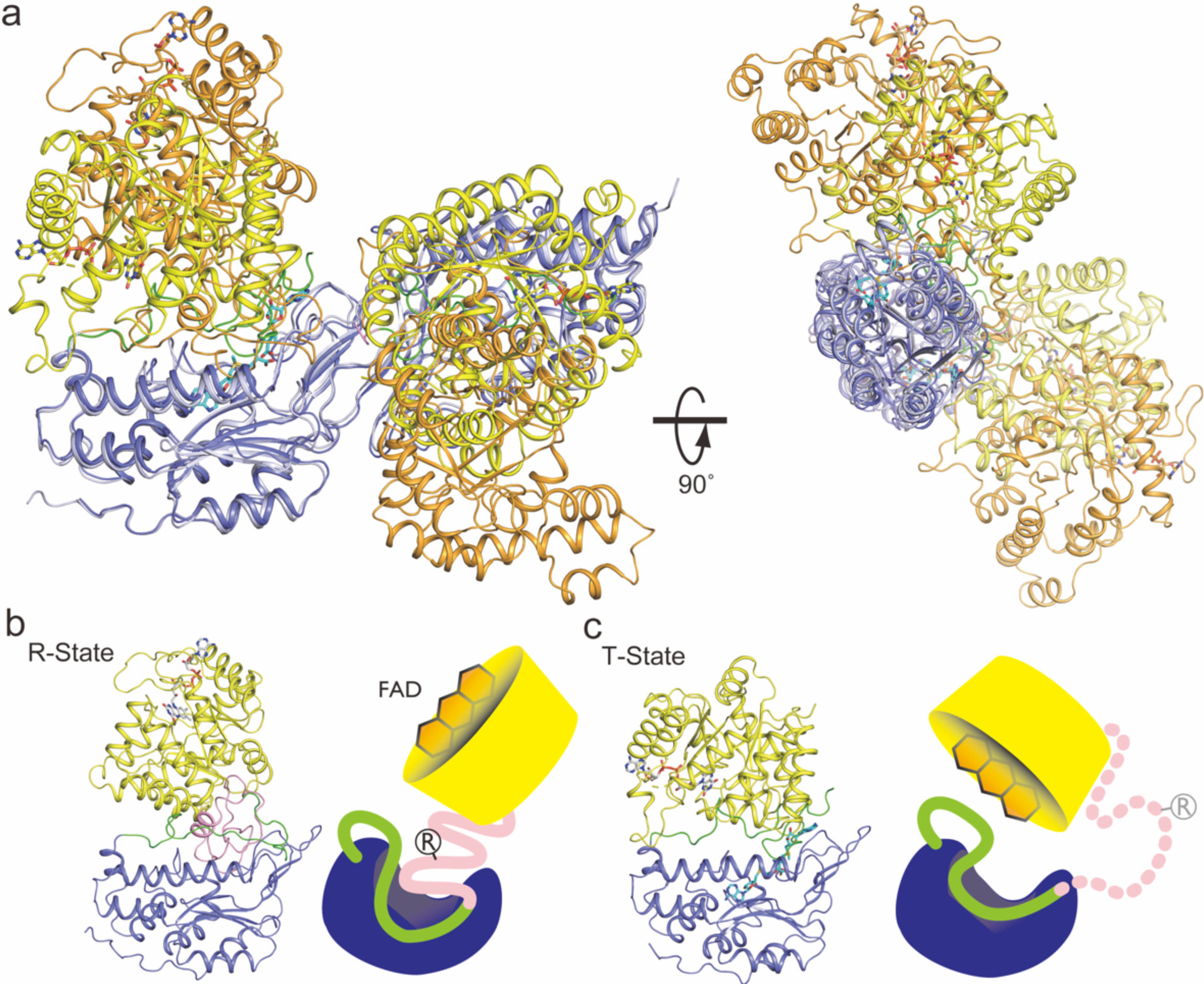
Comparison of *c*MTHFR in R- and T-state. **a** Dimer comparison of the R and T-states. The catalytic domain and regulatory domain are colored yellow and blue, respectively for the T-state, and light orange and cyan, respectively for the R-state. **b** Monomer in the R-state illustrated in **b** and monomer in the T-state in Panel **c** using the same color scheme as in Figure 3; these are also drawn in cartoon style. Since the structure in T-state lacks the N-terminal part of the linker (pink), the region is drawn as a dotted line in the cartoon. In the R-state, MTHFR accommodates the N-terminal pink region of the linker between the catalytic and regulatory domains. The C-terminal portion of the linker conceals the active site, resulting in substrates binding to the active site of MTHFR. The linker part is herein called in the inhibitory region. The circled R shows the Arg residue mutated to obtain the “T-state-lock” MTHFR, which is on the retractable region. The R and T MTHFR states are illustrated in a schematic mode.

The alignment between the two regulatory domains when compared is indistinguishable (RMSD ∼0.39 Å, Supplementary Fig. 9a), highlighting a conserved β-sandwich dimer interface mediated between adjacent regulatory domains in both configurations. Superposition of the catalytic domains likewise demonstrates that they are indistinguishable between the two different states (RMSD ∼0.53 Å, Supplementary Fig. 9b). However, in the T-state, the catalytic domains re-orient with respect to the regulatory domain, and MTHFR in this inactive state adopts a more compact conformation overall. Consequently, the transition between the R- to the T-state leads to an overall “contraction” of the global structure that shifts the catalytic domains (and their FAD centers) ∼23 Å closer to one another.

The catalytic domain transitions relative to the regulatory domain, from R- to the T-state, are primarily driven by the rearrangement of the linker. This flexible linker (aa293-370) consists of two regions an N-terminal retractable-hinge (aa293-348, pink, Figs. 4, 5, 6 and 7) and a C-terminal velcro-wedge (aa349-370, green, Fig. 4, 5, 6, and 7), so called because it forms a new interface between the catalytic and regulatory domains (velcro) and occludes access to FAD via Tyr361 (wedge). The linker is arguably where the most drastic structural changes are observed (Figs. 4b, 5a, and 6a). The active site, particularly the FAD cofactor, becomes buried within the catalytic domain due to the rearrangement of the linker, with the velcro-wedge region embedded between the regulatory and catalytic domains, forming a new interface. The N-terminal retractable-hinge region undergoes a helix-loop transition between the R- and T-states, serving as a hinge that guides and governs the observed global structural changes. The most prominent structural feature of the retractable-hinge region in the R-state is the presence of an α-helix (Fig. 4b, 5a, and 6a) (aa329-337), similar to that observed in *h*MTHFR (aa370-379); most of the retractable-hinge region could not be modeled (aa296-336) in the T-state, indicating increased flexibility and/or increased solvent exposure when compared to the R-state, where it was embedded between the catalytic and C-terminal velcro-wedge region. Our structures identified hotspots (near Arg315 in *c*MTHFR) in structural regions of known allosteric importance (Fig. 5). When the linker between the catalytic and regulatory domains is retracted, Arg357 (*h*MTHFR) interacts with Thr378 and Glu368 (Fig. 5a, b). Thus, loss of Arg357 in the patient mutation leads to destabilization of the retractable-hinge region. Arg357 would be a critical amino acid residue for stabilizing the R-state: near Arg357 there are four positively charged amino acid side chains (Lys356, Arg358, Arg363 and Arg377) that form a positively charged patch. The positively charged patch is conserved in vertebrate MTHFR (Fig. 5c) and *c*MTHFR (except for Lys356, Fig. 5c).

**Figure 5.**
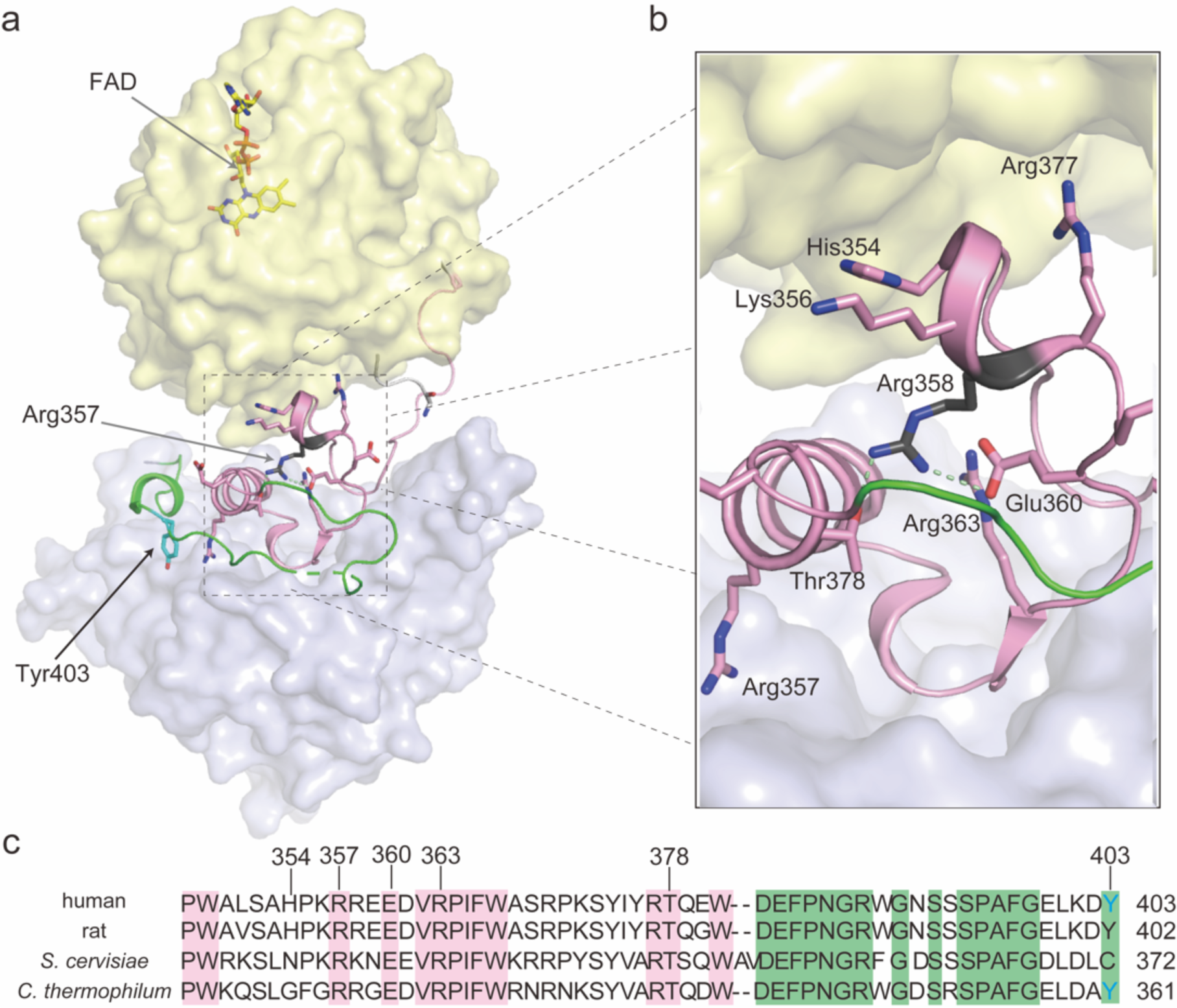
The role of the linker in interdomain interactions in the R-state. **a** The position of Arg357 in R-state. The Arg357Cys mutation is critical to destabilize the R-state or to stabilize the T-state because the mutation yields T-state-lock *h*MTHFR. **b** The retractable-hinge mediated linker interactions between domains, ostensibly crucial for the allosteric transition. Arg357 interacts with Thr368 and Glu360, which could stabilize the R-state, indicating that loss of the positively charged side chain by Cys mutation would destabilize the R-state. Moreover, there are five positively charged side chains, Lys356, Arg358, Arg377, and Arg363, found near Arg357. Given that the N-terminal phosphorylation site is close to the retractable-hinge region, the negatively charged phosphorylation sites would attract those positively charged side chains. **c** Amino acid alignment of MTHFR linker from *Homo sapiens*, *Rattus norvegicus*, *Saccharomyces cerevisiae*, and *Chaetomium thermophilum*. Conserved amino acids are highlighted in pink and green for the retractable-hinge and velcro-wedge regions, respectively.

**Figure 6.**
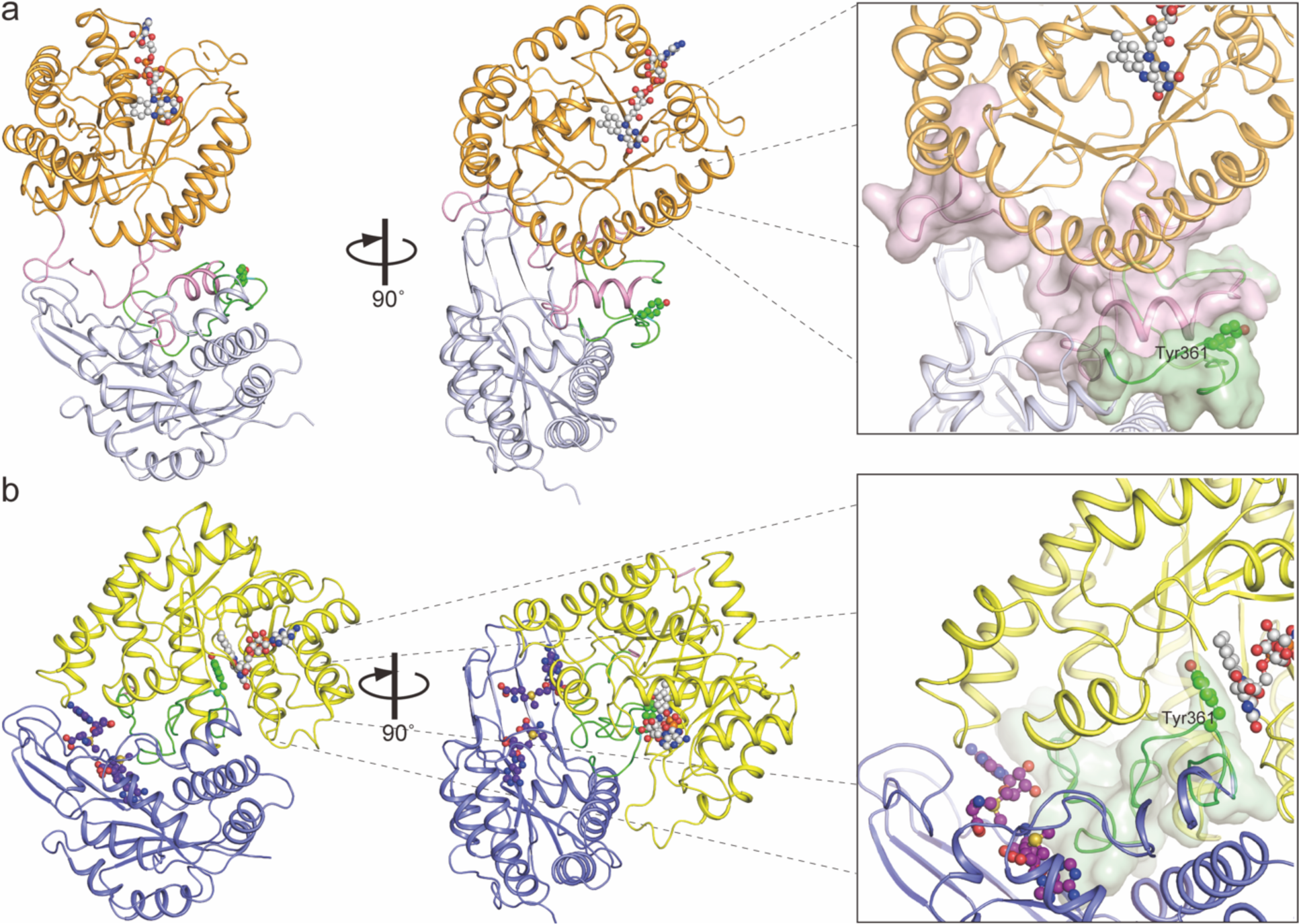
Linker region occludes FAD in MTHFR T-state. Structures of *c*MTHFR in R-state (**a**) and T-state (**b**) are shown using the same coloring scheme as in Figure 4. Tyr361, present on the C-terminal portion of the linker, occludes the *si*-face of FAD in the T-state, while that portion of the linker remains solvent-exposed in the R-state.

**Figure 7.**
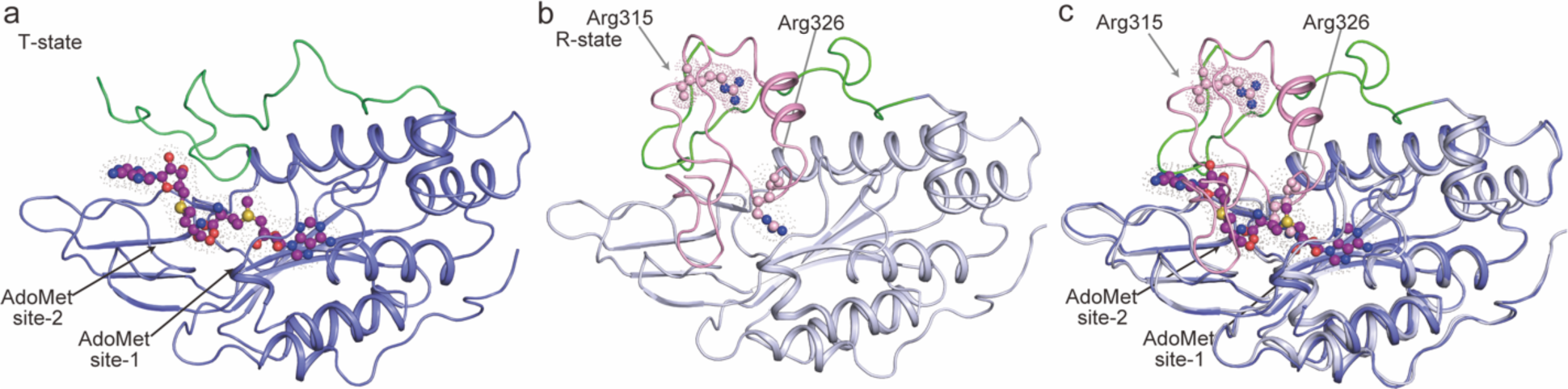
Effector molecule binding site comparison suggests a possible mechanism for how MTHFR transitions from the R to T-state is triggered. Structures of regulatory domain of *c*MTHFR in T-state (Panel **a**) and R-state (Panel **b**).Two AdoMet binding sites in T-state are surrounded by the regulatory domain, the inhibitory region of the linker, and the catalytic domain. The catalytic domain doesn’t contact with the regulatory domain in the R-state (Panel **b**) because of the presence of the retractable region of the linker. To demonstrate the effect of AdoMet-binding on MTHFR, the regulatory domain and the retractable region of the linker are superimposed (Panel **c**). AdoMet and AdoHcy share the AdoMet-site-1 (**d**). As AdoMet binds to the AdoMet site-1, the *S*-methyl moiety of AdoMet causes steric clash with Arg326 (**c**). The steric clash induces the reposition of the retractable region and disfavors the R-state. Repositioning of the retractable region reveals the AdoMet site-2 to facilitate the formation of the T-state.

The N-terminal linker region can be thought of as a hinge, whereby its extension causes a corresponding contraction/compacting of the catalytic domain towards the regulatory domain, yielding the T-state configuration. Indeed, this region occludes part of the allosteric site in the R-state (aa303-306, HALP). Additionally, the C-terminal velcro-wedge region serves to form part of the allosteric binding pocket (aa339-347) in the T-state structure, with aa395-409 of the regulatory domain acting as a lid over the allosteric site (Fig. 7). As such, the N-terminal linker region can be thought to act as a wedge in the R-state, whereby its occlusion of the secondary AdoMet binding site favors the R-state conformation.

The C-terminal velcro-wedge region, previously solvent exposed in the R-state, becomes completely embedded in a new interface between the catalytic and regulatory domains. Tyr361 on the velcro-wedge region repositions a dramatic ∼16 Å, forming a π-stacking interaction with FAD, directly occluding its *si-*face (Fig. 6b). The combined effect of these changes explains the catalytically inhibited nature of the T-state (Fig. 6b), where the Tyr361 and the C-terminal linker (Tyr finger) act as an autoinhibitory element, a form of intrasteric (active-site directed) regulation^23,24^. The movement of this loop/linker effectively serves to block access to the catalytic domain, trapping the enzyme in an inactive state while providing a closed pocket and stabilizing interaction with FAD that simultaneously explains the abolished catalytic activity *and* retention of the FAD cofactor observed with *c*MTHFR^R315A^; it also explains the altered UV-Vis spectrum observed upon AdoMet binding (Figs. 2c, d), which could thus be attributed to the altered FAD microenvironment^25^ in the T-state configuration. As such, the C-terminal linker region can, in turn, be thought to act as a lock/wedge in the T-state, whereby its interaction with the *si*-face of FAD, protrusion into the catalytic domain, and (re)positioning between the catalytic and regulatory domains serve to stabilize and favor the T-state.

### Binding of Two AdoMet molecules Promotes Drastic Linker Rearrangements

The regulatory domain was found to bind AdoMet, as expected for the T-state. Unexpectedly, two molecules of AdoMet were found per regulatory domain. One AdoMet molecule is found in the same site that AdoHcy occupies in the R-state (site-1), while a second AdoMet molecule is bound in a second adjacent site (site-2) that was occluded by residues of the retractable-hinge region (aa301-305) (Fig. 7, Supplementary Figs. 7 and 8) and whose pocket is in part lined by the velcro-wedge region (aa339-347) (Fig. 7a). Although the second AdoMet binding site is hidden in the R-state, the large conformational change that defines the T-state allows for the accommodation of *two* AdoMet molecules per monomer. AdoMet binding to site-1 and a steric clash with Arg326 on the retractable-hinge region or Ala408 (structurally analogous to Ala368 and Ala461 in *h*MTHFR, respectively) could precipitate the initial helix-loop transition, whose extension corresponds to movement of the catalytic domain. Previous work identified residues important to AdoMet binding, finding their sequence to be “s/t-Y-1-Y-x-T-Q-x-x-d/v-x-F-P-N-G-R-x-G-A-S.” in porcine MTHFR (aa373-SYIYRTQEWDEFPNGRWGNS-392 in *h*MTHFR, Fig. 5c). In the crystal structure of *c*MTHFR^R315A^, the peptide aa338-DWDEFPNGRWGDSRSPAFGELDA-360 occupies the corresponding position (Fig. 5c). While the AdoHcy-binding site (AdoMet site-1) is separated from this peptide, it is close to the second AdoMet-binding site (AdoMet site-2), suggesting that the previous study might have revealed the location of the second, cryptic AdoMet-binding pocket (AdoMet site-2)^26^. Another study identified residues important to AdoMet binding, where residues in AdoMet site-2 were surprisingly found to be important for AdoMet binding as a whole, despite them corresponding to the linker and not the regulatory domain^27^. The unveiling of a cryptic AdoMet site previously occupied by aa297-DRPLKHALPW-307 of the retractable-hinge region of the linker in the R-state to T-state transition helps explain this previously puzzling finding.

MTHFR is an allosteric enzyme that undergoes slow, hysteretic changes in activity in response to its physiological regulator, AdoMet (lag)^1,17^. This was initially ascribed to the timing of retraction/retention of the hinge region upon AdoMet binding to the R-state, where the triggered conformational change represents the rate-limiting step between the transition from the R-state to the T-state, followed by a slow “isomerization” towards the final, inhibited form (T-state). However, the authors did acknowledge the possibility that the biphasic nature of AdoMet-inhibition could be explained by *multiple* AdoMet binding events, an initial “burst” phase for the primary AdoMet binding event (R-state, site-1), and a secondary AdoMet binding event to an intermediary configuration that hastens the allosteric transition (to T-state, site-2)^17^. The key to understanding this biochemical data was the adoption of an effector-induced pathway model wherein binding of one AdoMet molecule to the R-state triggers a conformational change that favors and allows for binding of a second AdoMet molecule to this intermediary state, providing the committed step towards the T-state, with cooperative AdoMet binding^17^.

While the regulatory domain remains relatively static between both captured configurations, aa318-326 of the linker (retractable-hinge region) interacts with the dimer interface in the R-state, while aa296-336 overall becomes solvent-exposed and flexible in the T-state. The linker serves a dual role as a “sensor” and hinge, as the N-terminal retractable-hinge region (aa293-348) undergoes a helix-loop transition, extending and unwinding in the R- to T-state transition (Figs. 7 and 8). AdoMet binding to site-1 is the likely impetus for the initial structural changes that precipitates the transition, as site-2 is only revealed in the T-state (i.e. after AdoMet site-1 is occupied). This implies that the initial effector-mediated remodeling restructures the regulatory domain as well, unveiling the cryptic AdoMet site and adopting a conformation that favors the binding of a second AdoMet molecule.

**Figure 8.**
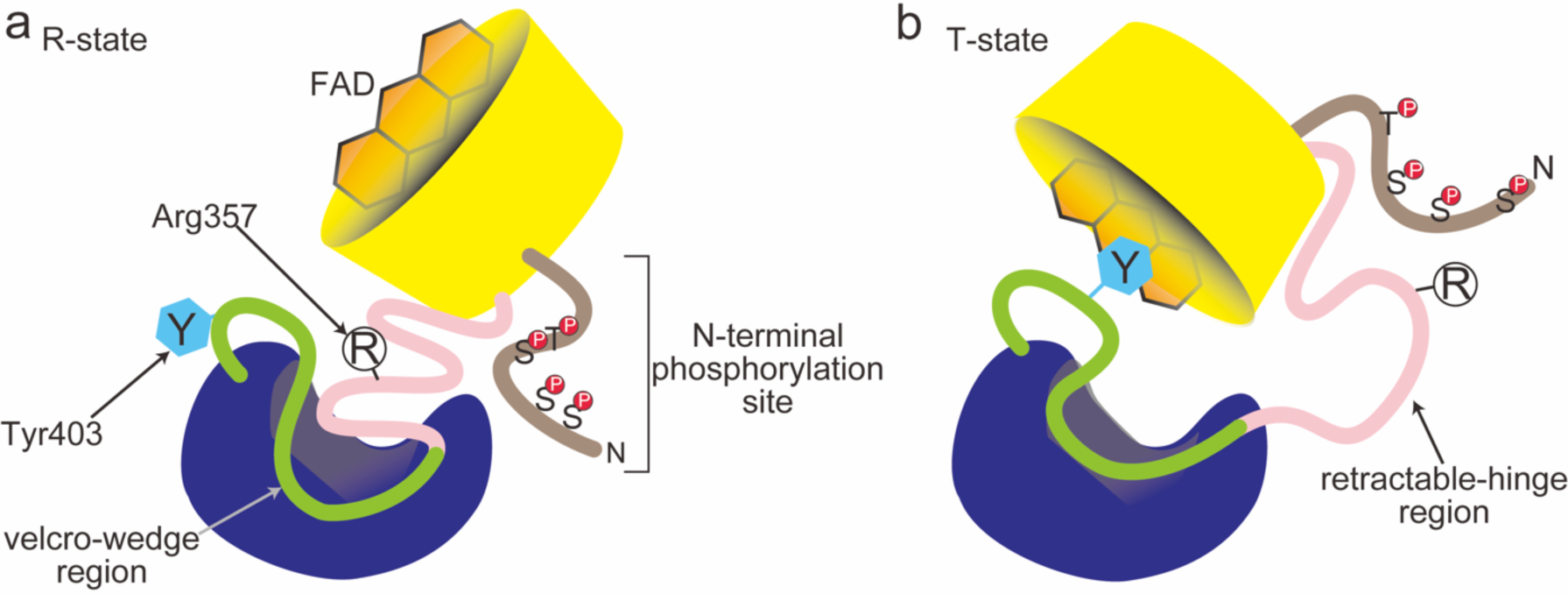
A proposed mechanism for how phosphorylation of *h*MTHFR enhances the AdoMet-sensitivity. **a** The N-terminal phosphorylation site (brown) is near the retractable region of the linker (pink) in the R-state. **b** In the T-state, the phosphorylation site and the retractable region may interact, although the retractable region would be flexible in solvent.

## Discussion

The regulation of eukaryotic MTHFR has proven to be multipronged and multifaceted, relying on phosphorylation status and effector-induced and stabilized conformational changes, all driven and achieved via the restructuring of its linker. Allostery via linker grants MTHFR a multimodal allosteric regulatory (cap)ability. The intricately layered role played by the linker, granted by its two disparate functional regions, indicates that MTHFR has evolved an original and, to our knowledge, unprecedented way of regulating its function via effector-mediated changes. Using a linker, eukaryotic MTHFR has shown that it can channel an initial chemical event, such as AdoMet binding, into a complex mechanical rearrangement whereby it orchestrates a drastic global contraction of the dimer, organizing a coupled rotation and translation of the catalytic domain (45° and ∼16 Å, respectively). In addition, it simultaneously reveals a secondary cryptic AdoMet site and occluding access to the *si*-face of FAD. Most astonishingly, a portion of the linker ends up being buried between the regulatory and catalytic domains, though it still participates in AdoMet site-2, FAD, and T-state stabilization. The transition from the R- to T-state involves a drastic decrease in the surface area of the overall structure, with a concurrent contraction reminiscent of hydrophobic collapse. The velcro-wedge region of the linker is solvent exposed in the R-state but becomes completely buried in the T-state; the opposite is true for the retractable-hinge region, where its environment is hydrophobic in the R-state but becomes solvent exposed in the T-state.

It is worth mentioning that phosphorylation itself must be considered to play an important role, especially in *h*MTHFR, where it has been shown that phosphorylation of predominantly N-terminal residues increases AdoMet-dependent allosteric inhibition. The retractable-hinge region of the linker is flexible, particularly in the T-state, and as such it can presumably interact with the phosphorylated amino acid side chains of the N-terminal (Fig. 8). The phosphorylation status of MTHFR adds yet another layer to the mechanism by which effector-mediated structural changes are channeled via the linker.

This effector-mediated allosteric conformational change is wholly unexpected, as is the unveiling of a cryptic secondary AdoMet binding site. AdoMet site-2 is only unveiled after AdoMet site-1 is occupied and the conformational rearrangement has begun. AdoMet site-2’s direct proximity and interaction with the velcro-wedge region of the linker (F342) and the regulatory domain (W447) (Supplementary Fig. 7), along with its more solvent exposed nature relative to site-1, make it a prime candidate site to initiate the reverse transition (T- to R-state). Therefore, signaling that AdoMet/AdoHcy concentrations in the cell require decreased flux into the folate cycle could be committing the enzyme to T- to the R-state transition. It has been shown that NADPH binding precipitates the reverse transition (T- to R-state), eliciting the opposite effect of AdoMet binding^17^. However, how the T-state, once attained, can transition to and rearrange back to the R-state is not clear. In the R-state, when AdoMet binds to site-1, it is proximal to and clashes with the Arg326 retractable-hinge region of the linker, providing a direct method via which the initial AdoMet binding event to site-1 can trigger the observed conformational changes (Fig. 7c).

The linker region likely has additional roles to those mentioned above. The flexibility of the linker in the T-state in our experimental structures and computational models hints that it is also crucial in the T- to R-state transition. As it stands, while potential mechanisms to transition from the T- to the R-state are postulated, such as dephosphorylation of PTM residues, AdoHcy-binding and/or AdoMet-unbinding, and NADPH-binding, these remain solely theoretical, and further work must be done to ascertain what governs T- to R-state transition (“reactivation”). While it is prudent to assume that the linker will play a major role, the initial event that triggers the reconfiguration and the interactions the retractable-hinge and velcro-wedge regions could form are unclear.

The outsized importance of MTHFR as the convergence point and rate-limiting enzyme in one-carbon metabolism has forced Nature to adopt two common structural motifs (TIM barrel, β-sheet) and use unheralded features, in this case, a linker, to achieve a novel form of allostery. Through a linker region, MTHFR can govern one-carbon metabolism flux, sense and react to cellular conditions (AdoMet/AdoHcy concentration and ratio), and use ∼80 amino acids to sense, block, protect, and trigger dramatic global conformational changes. Though already complex, the structural and mechanistic details regarding MTHFR allostery are only just being described. Whether MTHFR’s unique take on allostery via the use of its linker is more common than previously appreciated remains unclear. The use of two allosteric inhibitor sites, one cryptic and only exposed after the initial inhibitor binding event, represents a unique remodeling of an allosteric site; the unexpected stoichiometry of two AdoMet molecules bound per monomer indicates that each site plays a differing role in the allosteric regulation and subsequent structural transition between R- and T-states and that Nature has strategically tuned its ability to sense and respond to the AdoMet/AdoHcy ratio present in the cell, gating one carbon flux accordingly.

## Methods

### Preparation of Recombinant *h*MTHFR

Preparation of recombinant *h*MTHFR was performed according to protocols described previously^14,20^. Both the wild-type variant and the Arg357Cys mutant were obtained via the baculovirus-insect cell expression system. Generation of a donor vector for the Arg357Cys mutant was performed using the QuikChange mutagenesis kit (Agilent). Briefly, recombinant *h*MTHFR obtained from Sf9 cells was resuspended in 50 mM potassium phosphate buffer (KPB), pH 7.2, 0.1 M NaCl, and 1% Triton X-100 (8 mL per 1 g of pellet). The resuspended cell pellet was lysed via sonication (4 °C, 10 min). The crude lysate was centrifuged (20 min, 5,000 x g, 4 °C), decanting the supernatant to remove any cellular debris/pellet. To this supernatant, 2 grams of DEAE cellulose (DE-52, Whatman) was added and stirred in an ice-water bath. The resulting gel slurry was then packed into an empty 2.5 cm diameter column. After a wash with 50 mM KPB, pH 7.2, and 0.1 M NaCl (50 mL), the protein was eluted in bulk with 50 mM KPB, pH 7.2, and 0.3 M NaCl (20 mL). The eluate was then applied onto a Ni-affinity column (His-Trap Chelating HP, Cytvia/GE, 1 mL) pre-equilibrated with 50 mM KPB, pH 7.2, 0.3 M NaCl, and 0.05 M imidazole. Following a wash (50 mM KPB, pH 7.2, 0.3 M NaCl, and 0.1 M imidazole), the protein was eluted in bulk (50 mM KPB, pH 7.2, 0.3 M NaCl, and 0.3 M imidazole). The eluted fraction was dialyzed at 4 °C overnight (50 mM KPB, pH 7.2). The resulting dialysate was concentrated and stored at −80 °C.

### Preparation of Recombinant *c*MTHFR

The cDNA encoding MTHFR was sourced from *Chaetomium thermophilum var. thermophilum* DSM 1495 [GenBank accession code XP_006693807] (NCBI Gene Locus tag CTHT_0033700). *C. thermophilum* possesses two MTHFR genes with locus tags CTHT_0065570 and CTHT_0033700. The latter was selected for its greater sequence similarity to the *Saccharomyces cerevisiae* Met13 gene^28^, whose activity is modulated by AdoMet, and was synthesized by GeneArt (Invitrogen). The wild-type gene, originally cloned in a pMA vector, was subcloned in a pMSCG7 vector using ligation-independent cloning (LIC). The expression vector was designated as pMCSG7(*c*MTHR^wt^). To express the R315A mutant and E21Q, L393M, V516F triple mutant, site-directed mutagenesis was performed using the QuikChange mutagenesis kit (Agilent) to construct pMCSG7(*c*MTHR^R315A^) and pMCSG7(*c*MTHFR^E21Q, L393M, V516F^). *BL21star(DE3)* was used for protein expression. *E. coli* transformed with either pMCSG7(*c*MTHR^wt^), pMCSG7(*c*MTHR^R315A^), or pMCSG7(*c*MTHFR^E21Q, L393M, V516F^) was propagated at 37 °C in Luria Broth containing 50 µg/mL ampicillin, and protein overexpression was induced via auto-induction^29,30^. Cells were grown at 30 °C overnight before harvesting via centrifugation and stored at −80 °C. The harvested cell pellet was resuspended in 50 mM KPB, pH 7.4 (4 mL per 1 g of pellet), to which lysozyme (0.1 mg/mL) and PMSF (1 mM) were added. The resuspended cell pellet was lysed via sonication (4 °C, 5 s on, 5 s off, 5 min total). The crude lysate was centrifuged (45 min, 20,000 x g, 4°C), decanting the supernatant to remove any cellular debris/pellet. The crude supernatant was collected and loaded onto a Ni-affinity column (His-trap Chelating HP, Cytvia/GE, 5 mL) pre-equilibrated with 50 mM KPB, pH 7.4, and 20 mM imidazole. The column was washed using the equilibration buffer (20 mM imidazole), then 80 mM imidazole, and the protein was eluted in bulk with a buffer consisting of 50 mM KPB, pH 7.4, and 250 mM imidazole. Protein-containing fractions as judged by SDS-PAGE analysis were collected and subjected to a TEV digest, with dialysis at 4 °C overnight (50 mM KPB, pH 7.4). The dialysate was loaded onto a pre-equilibrated HiLoad 16/600 Superdex 200 pg gel filtration column (SEC buffer, 25 mM Tris, pH 7.4, 0.1 M KCl, and 1 mM TCEP). Fractions containing the desired protein, as judged by SDS–PAGE, were pooled and concentrated via centrifugation in 25 mM Tris, pH 7.4, 50 mM KCl, 1 mM TCEP to yield purified *c*MTHFR^wt^/*c*MTHFR^E21Q, L393M, V516F^ (∼20 mg/mL) or *c*MTHFR^R315A^ (∼50 mg/mL), which was stored at 4 °C or flash-frozen for long-term storage at −80 °C.

### Biochemical analysis of *h*MTHFR patient mutations

The recombinant *h*MTHFR^R357C^ mutant was produced using the baculovirus-insect cell expression system^20^. Activity for the cell extract expressing histidine-tagged *h*MTHFR^R357C^ was assessed using the NADPH:mendione oxidoreductase assay. FAD release from *h*MTHFR was measured according to our previous method^20^, with minor modifications. A spectrofluorophotometer RF-5300PC (Shimadzu) with a cell-temperature controller was used. The excitation and emission wavelengths were set at 390 nm and 525 nm, respectively. Concentrated *h*MTHFR was diluted directly into pre-warmed 50 mM KPB, pH 7.2 (3 mL) at 46 °C to a final concentration of 100 nM, and the fluorescence intensity from released FAD was monitored for 10 min.

### Enzyme assay and UV-Vis spectroscopy of *c*MTHFR

Enzyme assays using *c*MTHFR were performed as described previously^2^, with some minor modifications. NADPH:menadione oxidoreductase activity was measured using a Cary 100 Bio spectrophotometer (Agilent Technologies, Inc). The reaction mixture without menadione was prepared at room temperature, containing 50 nM of *c*MTHFR with 100 μM NADPH or NADH in 50 mM KPB, pH 7.2. Reactions were initiated by adding concentrated menadione solution (100 μL per mL) to the cuvette. The consumption of NADPH (or NADH) was monitored via absorbance at 343 nm at room temperature. The change in concentration of NAD(P)H was determined using the extinction coefficient of oxidized NAD(P)H of 6,220 M^−1^ cm^−1^ at 343 nm. Once NADPH preference was established, the effect of *S*-adenosylmethionine (AdoMet) on activity was assessed by adding 100 μM of AdoMet to the reaction mixture before menadione addition. Similarly, to test the effect of FAD on activity, 2 μM of FAD was added to the reaction mixture before menadione addition. Comparison of the UV-VIS spectra of *c*MTHFR, monitoring the changes at 450 nm associated with FAD, were assessed between *c*MTHFR^wt^, *c*MTHFR^wt^ in the presence of 100 μM of AdoMet, and *c*MTHFR^R315C^. To determine the changes in absorbance at 450 nm as a function of AdoMet concentration, spectra were recorded using 50 nM of *c*MTHFR^wt^ and 0–200 µM of AdoMet.

### Surface lysine methylation

For *c*MTHFR crystallization in the R-state, purified protein (*c*MTHFR^E21Q, L393M, V516F^) underwent treatment with formaldehyde and dimethylamine-borane complex (Sigma-Aldrich) for surface lysine methylation^31,32^. After lysine methylation, the protein was loaded onto a pre-equilibrated HiLoad 16/600 Superdex 200 pg gel filtration column (SEC buffer, 25 mM Tris, pH 7.4, 0.1 M KCl, and 1 mM TCEP). Fractions containing the desired protein, as judged by SDS–PAGE, were pooled and concentrated via centrifugation in 25 mM Tris, pH 7.4, 50 mM KCl, 1 mM TCEP to yield surface lysine-methylated *c*MTHFR^E21Q, L393M, V516F^ (∼20 mg/mL) which was stored at 4 °C or flash-frozen for long-term storage at −80 °C.

### Post-translational modification of recombinant *h*MTHFR

Phosphorylation sites were analyzed according to previous reports^33,34^. Purified His-tagged recombinant wildtype *h*MTHFR (∼130 µM, 120 µL) was treated with 10 mM DTT for 30 min at room temperature, followed by iodoacetamide (50 mM final concentration) for an additional 30 min at room temperature. Trypsin (20 µg, Pierce Trypsin protease, MS-Grade) was used for digestion and the mixture was incubated at 37 °C overnight. The reaction was quenched by adding TFA (final concentration, 0.5% v/v). Phosphopeptides were enriched using a TiO_2_ column (GL Science) and then concentrated using a C_18_ tip column (ZipTip, Millipore). Phosphopeptide-enriched tryptic fragments were analyzed using a Thermo Scientific Orbitrap Fusion Lumos Tribrid MS coupled with the UltiMate 3000 RSLCnano liquid chromatography system (University of Michigan, Department of Chemistry). The proteomics data analysis was done using Proteome Discoverer (Thermo Scientific).

### Crystallization of *c*MTHFR

Crystals of the inhibited form (T-state) of *c*MTHFR were grown as follows: Fifty mg/ml (∼210 µM) *c*MTHFR^R315A^ was prepared in 50 mM KPB, pH 7.4 containing 500 µM AdoMet and 250 µM FAD. The protein solution was mixed in a 1:1 ratio with the reservoir solution (0.1 M sodium acetate, pH 4.8, 0.2 M ammonium sulfate, 6% polyethylene glycol monomethyl ether 2,000). Crystals were grown at 20 °C using the hanging-drop vapor diffusion method. Crystals of *c*MTHFR in the active form (R-state) were obtained using a *c*MTHFR Glu21Gln, Leu393Met, Val516Phe mutant (*c*MTHFR^E21Q, L393M, V516F^), whose surface lysine residues were modified via reductive methylation. Twenty mg/mL of *c*MTHFR^E21Q, L393M, V516F^ was prepared in 25 mM Tris, pH 7.4 containing 50 mM KCl, 500 µM FAD, and 1 mM TCEP. The protein solution was mixed in a 1:1 ratio with the reservoir solution (0.1 M HEPES, pH 7.5, 0.1 mM potassium chloride, 20 mM magnesium chloride, 22% poly(acrylic acid sodium salt) 5,100). Crystals were grown at 20 °C using the sitting-drop vapor diffusion technique. Crystals were briefly transferred to a cryo-protectant solution (the reservoir solution containing 20% glycerol), 1 min for T-state crystals, and 30 sec for R-state crystals, before harvesting and flash freezing in liquid nitrogen.

Data collection and processing statistics are summarized in Supplementary Table 1. Data for *c*MTHFR^R315A^ were indexed to space group *P*22_1_2_1_ (unit-cell parameters *a* = 130.66, *b* = 149.95, *c* = 171.06 Å) with four molecules in the asymmetric unit (Matthew’s coefficient VM = 2.99 Å^3^ Da^−1^, 59% solvent content). Data for *c*MTHFR^E21Q, L393M, V516F^ were indexed to space group *P*2_1_2_1_2_1_ (unit-cell parameters *a* = 117.97, *b* = 151.38, *c* = 188.05 Å) with four molecules in the asymmetric unit (Matthew’s coefficient VM = 3.00 Å^3^ Da^−1^, 59% solvent content).

### Data collection and refinement

X-ray data sets were collected at 100 K on LS-CAT beamline 21-ID-D at the Advanced Photon Source, Argonne National Laboratory (Argonne, IL). Data sets were processed using xia2/DIALS^35^. Initial phases for *c*MTHFR^R315A^ were obtained using Phaser^36^. The catalytic domain of *h*MTHFR (6FCX) and the regulatory domain of *h*MTHFR (6FCX) without ligands were used as an N-terminal and C-terminal rigid body, respectively, as search models for molecular replacement. Iterative model building and corrections were performed manually using Coot^37^ following molecular replacement, with the loop region being placed manually as a poly-alanine chain, and subsequent structure refinement was performed with CCP4 Refmac5^38^. PDB-REDO^39^ was used to assess the model quality in between refinements and to fix any rotamer and density fit outliers automatically. The model quality was evaluated using MolProbity^40^. For *c*MTHFR^E21Q, L393M, V516F^, initial phases were obtained via molecular replacement using MRBUMP^41^. The search model used was an AlphaFold model of an MTHFR homolog (UniProt A0A175W7M4). Iterative model building and corrections were performed manually using Coot^37^ following molecular replacement and subsequent structure refinement was performed with CCP4 Refmac5^38^. Initial refinement was conducted using BUSTER^42^ to rapidly fix Ramachandran, rotamer, and density fit outliers, refining to convergence and adding water molecules in the final automated round of refinement. Phenix eLBOW^43^ was used to generate the initial ligand restraints using ligand ID “FAD”. Phenix LigandFit^44^ was used to provide initial fits and placements of the FAD ligands. PDB-REDO^39^ was used to assess the model quality in between refinements, to fix any rotamer and density fit outliers, and to optimize the ligand geometry and density fit automatically. Figures showing crystal structures were generated in PyMOL^45^.

## Acknowledgements

The authors acknowledge the LS-CAT beamlines at the Advanced Photon Source for beamtime. This work was funded by Rackham Merit Fellowship (JM), National Science Foundation (CAREER 194517 to MK), and supported by Special Coordination Funds for Promoting Science and Technology from the Ministry of Education, Culture, Sports, Science and Technology through the Japan Science and Technology Agency (KY).

## Statistical Analysis and Reproducibility

Unless otherwise stated, at least three independent experiments were conducted for the functional assays. All attempts at replication were successful. The representative results are displayed in Figure 2a-d, and for Supplementary Figure 3b and c. Analysis and curve-fitting was performed using Prism 9.5.1 (Figure 2) and Kaleidagraph 4.1.4 (Supplementary Figure 3b and c).

## Data Availability

The structure coordinates and structure factors reported in this study have been deposited in the Protein Data Bank under accession codes 8UY1 (*c*MTHFR^E21Q, L393M, V516F^) and 8UY1 (*c*MTHFR^R315A^). PDB codes of previously published structures used in this study are 6FCX. All other data are available from the corresponding authors upon request. Source data are provided in this paper.

