## Supplementary Information for "Structural basis of *S*-adenosylmethionine-dependent Allosteric Transition from Active to Inactive States in Methylenetetrahydrofolate Reductase"

**1 and 2.** Identification of phosphorylation sites of recombinant wildtype *hMTHFR*

**3.** *hMTHFR* Arg357Cys patient mutation

**4.** FAD binding site in *cMTHFR*, R-state

**5.** Limited proteolysis of *cMTHFR* in the presence or absence of AdoMet

**6.** FAD and Tyr361 interaction in T-state

**7.** AdoMet binding sites in *cMTHFR*, T-State

**8.** Electron density around AdoMet allosteric inhibitors in *cMTHFR*, T-State

**9.** Structural rigidity of *cMTHFR* domain in R and T states

**Supplementary Table 1.** X-Ray Data Collection and Refinement Statistics

**Supplementary Table 2.** Bacterial and insect strains, plasmids, and synthetic oligonucleotides used in this study

**References**

**1 and 2**. **Identification of phosphorylation sites of recombinant wildtype *h*MTHFR**

Recombinant *h*MTHFR expressed in insect (Sf9) cells is highly phosphorylated; the phosphoryl groups can be removed by phosphatase treatment[1,2]. In the present study, we found 11 phosphorylation sites in *h*MTHFR. Although 16 phosphorylation sites were previously reported[2], including non-surface exposed sites such as Tyr90 in the catalytic domain, all amino acid residues identified by our analysis were exposed to solvent based on the *h*MTHFR structure (PDB, 6FCX)[2]. A summary of phosphorylation sites in recombinant wild-type *h*MTHFR in previously reported and our analyses is shown (Fig. 1). There are eight overlapping residues in the identified phosphorylation sites. Of particular significance are the seven residues within this overlap that populate the N-terminal Ser/Thr rich region, namely, Ser21, Ser23, Ser25, Ser26, Ser29, Ser30, and Thr34. The latter is thought to be the priming position for post-translational modification of *h*MTHFR by Pro-directed kinase(s). An additional phosphorylation site, Ser394, is located within the linker region. Kinases, including CDK1/cyclin B[3], polo-like kinase 1[4], DYRK1A/2[5], and GSK3A/B[5], have been proposed to participate in the post-translational modification of *h*MTHFR. Our LC-MS analysis revealed 11 phosphorylation sites in recombinant *h*MTHFR. More than two trypsin-digested phosphopeptide fragments were used to confirm these modifications required the use of the higher-energy collisional dissociation (HCD)-induced fragmentation mass spectra, facilitated phosphopeptide identification. Typical MSMS data are shown (Fig. 2).

AdoMet acts a more conspicuous inhibitory influence on the MTHFR activity of fully modified *h*MTHFR in contrast to phosphatase-treated *h*MTHFR at reduced concentrations. Furthermore, *h*MTHFR$^{T34A}$ also manifests in reduced responsiveness to AdoMet, possibly due to the Thr34 to Ala substitution impeding sequential post-translational modifications. This observation posits an association between the post-translational modification of *h*MTHFR and the allosteric regulation and transition of the enzyme. Moreover, in light of the attenuated sensitivity to AdoMet observed in the N-terminal truncated *h*MTHFR[2], it becomes apparent that the heavily phosphorylated N-terminal region assumes a pivotal role in the allosteric transition of *h*MTHFR to the T-state.

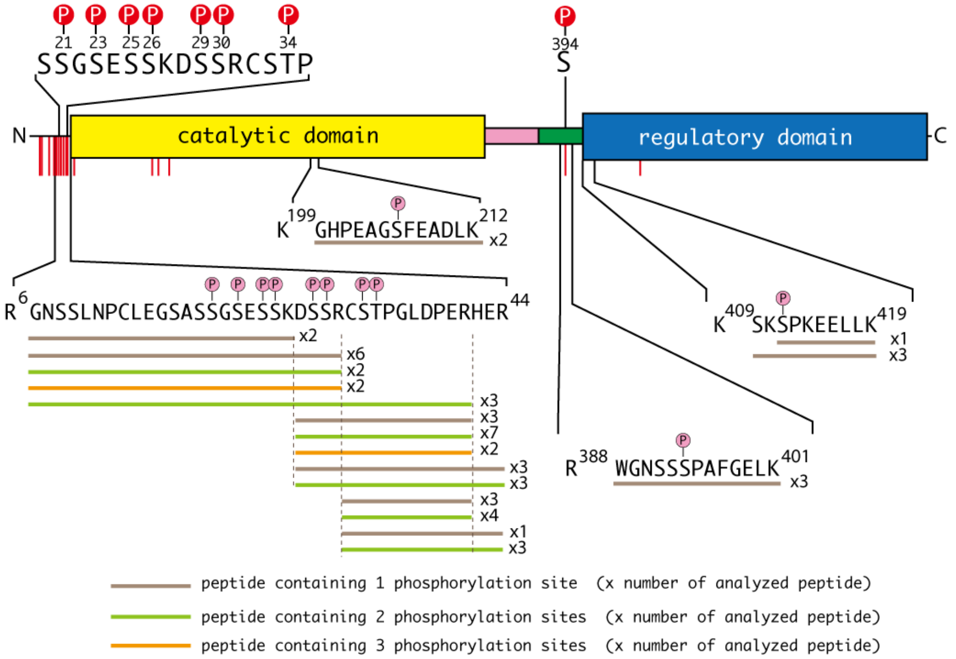


**Supplementary Figure 1. Schematic representation of the phosphorylation sites in recombinant *h*MTHFR.** The domain structure of *h*MTHFR is shown in a systematic mode. The catalytic and regulatory domains are shown in yellow and blue, respectively. The linker connecting the catalytic and regulatory domains is colored pink and green, denoting the "retractable region" and "inverted cap-for-active-site", respectively (see the main text for a detailed explanation). The upper part of the schematic domain illustration shows eight phosphorylated residues that are consistent in both the previous report and the present study. These residues are marked with red circled "P" letters. The previously reported phosphorylation sites are indicated by red vertical lines[2]. The phosphorylation sites identified in this study are described in the lower part. Tryptic fragments of *h*MTHFR were analyzed by LC-MS and phosphorylated amino acid residues identified in this study are marked with pink circled "P" symbols. The horizontal lines below the amino acid sequences represent the numerical distribution of the MS/MS fragments analyzed. Each color within these lines indicates the different amounts of phosphorylated amino acid residue(s) in a tryptic peptide, a classification explained in the lower section.

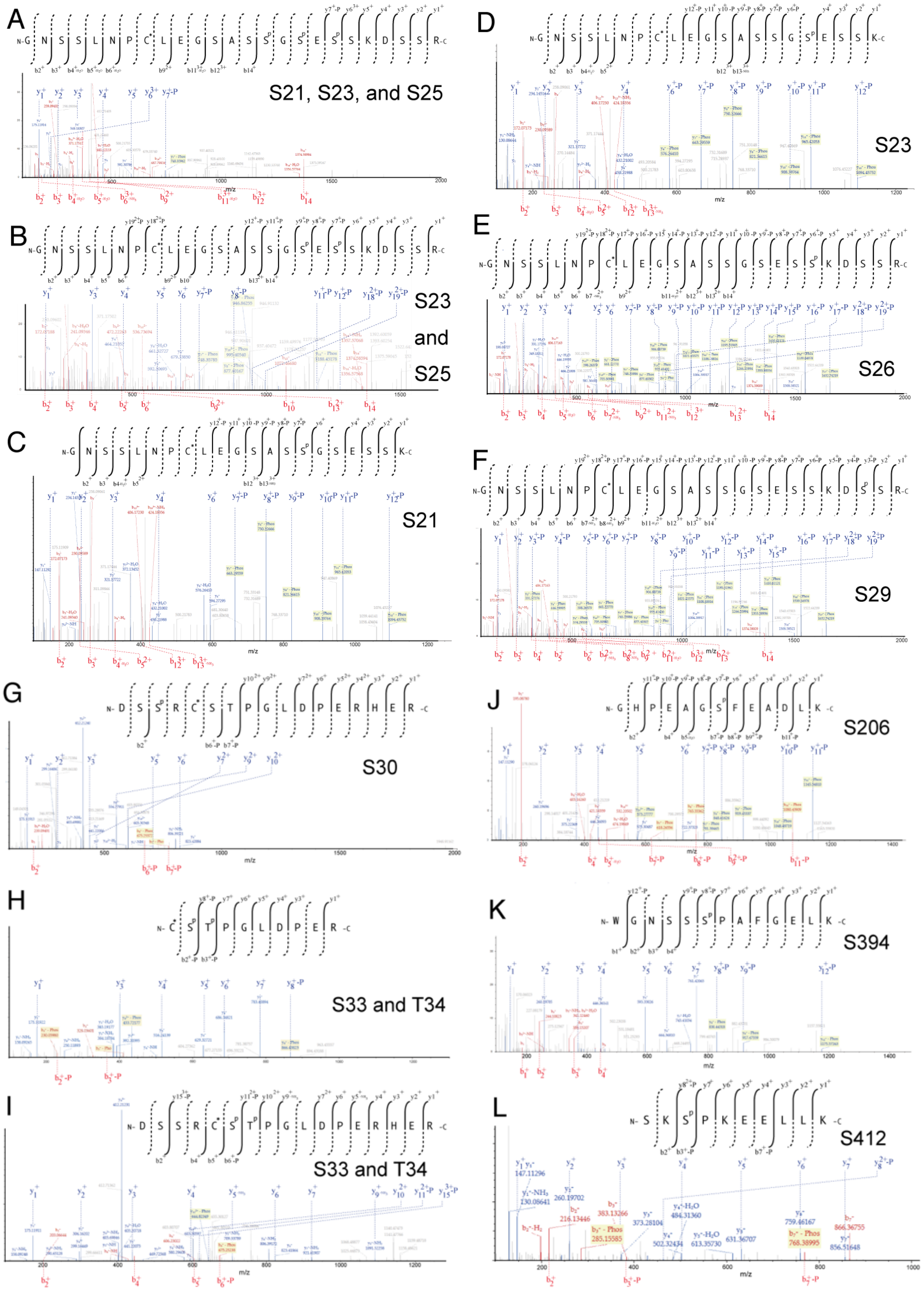

**Supplementary Figure 2. Identification of phosphorylation sites of recombinant *h*MTHFR by mass spectrometry.** (**a-l**) LC-MS/MS data showing Higher-energy collisional dissociation (HCD) -induced fragmentation mass spectra identifying eleven phosphopeptides. Observed b-ions are shown in red, whereas y-ions are shown in blue. **a** MS/MS spectrum of precursor m/z 914.00164 Da (+3) and MH + 2739.99038 Da, of the tryptic phosphopeptide GNSSLNP(*C)LEGSAS(pS)G(pS)E(pS)SKDSSR; where (*C) indicates carbamidomethylated Cys and (pS) represents for phosphorylated Ser. **b** MS/MS spectrum of precursor m/z 887.34474 (+3) and MH + 2660.01966 Da, of the tryptic phosphopeptide GNSSLNP(*C)LEGSASSG(pS)E(pS)SKDSSR. **c** MS/MS spectrum of precursor m/z 712.29251 (+3) and MH + 2134.86297 Da, of phosphopeptide GNSSLNP(*C)LEGSAS(pS)GSESSK. **d** MS/MS spectrum of precursor m/z 712.29251 (+3) and MH + 2134.86297 Da, of the tryptic phosphopeptide GNSSLNP(*C)LEGSASSG(pS)ESSK. **e** MS/MS spectrum of precursor m/z 860.68966 (+3) and MH + 2580.05444 Da, of the tryptic phosphopeptide GNSSLNP(*C)LEGSASSGSES(pS)KDSSR. **f** MS/MS spectrum of precursor m/z 860.68966 (+3) and MH + 2580.05444 Da, of the tryptic phosphopeptide GNSSLNP(*C)LEGSASSGSESSKD(pS)SR. **g** MS/MS spectrum of precursor m/z 552.89477 (+3) and MH + 1656.66977 Da, of the tryptic phosphopeptide DS(pS)R(*C)STPGLDPER. **h** MS/MS spectrum of precursor m/z 646.22458 (+3) and MH + 1291.44189 Da, of the tryptic phosphopeptide (*C)(pS)(pT)PGLDPER. **i** MS/MS spectrum of precursor m/z 540.46488 (+3) and MH + 2158.83768 Da, of the tryptic phosphopeptide DSSR(*C)(pS)(pT)PGLDPERHER. **j** MS/MS spectrum of precursor m/z 719.30610 (+2) and MH + 1437.60492 Da, of the tryptic phosphopeptide GHPEAG(pS)FEADLK. **k** MS/MS spectrum of precursor m/z 730.31791 (+2) and MH + 1459.62854 Da, of tryptic phosphopeptide WGNSS(pS)PAFGELK. **l** MS/MS spectrum of precursor m/z 413.55088 (+3) and MH + 1238.63807 Da, of tryptic phosphopeptide SK(pS)PKEELLK.

#### 3. *h*MTHFR Arg357Cys patient mutation

Over 100 mutations have been reported in the human MTHFR gene[6]. The Arg357Cys mutation in the MTHFR deficient patients results from the rare 1081C>T gene mutation[6,7,8]. The numbering of nucleotide and amino acid residues follows the nomenclature of the previous report[7]. The 1081C>T gene mutation is a rare mutation; it has been found in nine alleles in two families[6]. Patients with 1081C>T have low MTHFR activities, 5~27% of controls; thus, most of them are in severe (usually the activity range is 0%-20% of controls MTHFR deficiency[9,10].

The recombinant histidine-tagged *h*MTHFR Arg357Cys mutant was produced by a previously reported method employing the baculovirus-insect cell expression system[11]. The Arg357Cys mutant is chiefly found in the soluble fraction of the protein (Fig. 3a). The Arg357Cys mutant showed a slower mobility on SDS-PAGE in compared to the Thr34Ala mutant, which had no post-translational modifications[11]. Rather, it migrates with similar mobility as the wild-type enzyme. Therefore, it is likely that the mutant undergoes a post-translational modification, such as phosphorylation[11].

However, the cell extract expressing the *h*MTHFR Arg357Cys mutant showed only ~8% NADPH-menadione oxidoreductase activity compared to that expressing wild-type *h*MTHFR, and it was only slightly higher than the endogenous NADPH oxidase activity in the Sf9 cell. It is generally known that the NADPH-menadione oxidoreductase assay is not suitable for estimating recombinant MTHFR enzyme activity in cell lysate. However, our expression system allowed us to roughly estimate the recombinant MTHFR enzyme activity in the cell lysate due to the high level of the protein production. Routinely, NADPH-menadione oxidoreductase activity in uninfected Sf9 cells, a negative control for non-MTHFR-related activity, is less than ≈5% of that in baculovirus-infected Sf9 cells expressing wild-type *h*MTHFR. The high-level expression of *h*MTHFR facilitated the acquisition of purified recombinant mutants. Despite the use of the purified enzyme, the mutant exhibited an extremely low turnover number (Fig. 3b). Due to this low activity, the determination of NADPH affinity and AdoMet inhibition could not be determined.

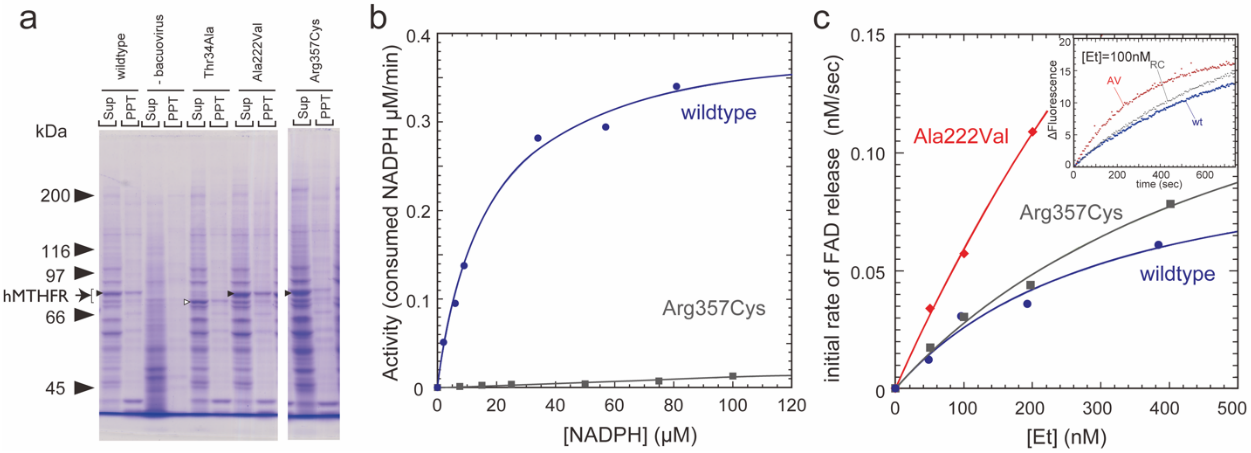


**Supplementary Figure 3. Heterologous expression, NADPH-menadione oxidoreductase activity, and FAD release of the Arg357Cys mutation. a** Insect cells, infected with baculovirus harboring *h*MTHFR genes, were lysed using a lysis buffer (10 mM Tris-HCl at pH 7.2, 2 mM ethylenediaminetetraacetic acid, 0.1 M NaCl, 50 mM sodium fluoride, 1 mM dithiothreitol, 1 mM phenylmethylsulfonyl fluoride, and 1% NP-40) on ice. The resulting cell lysate was subjected to centrifugation at 4˚C to obtain soluble (supernatant, Sup) and insoluble (precipitate, PPT) fractions, both of which were subsequently analyzed by SDS-PAGE. The gel was stained with Coomassie brilliant blue, with arrowheads indicating *h*MTHFR cDNA products. The Thr34Ala mutant (indicated by a white arrowhead) exhibited faster migration, due to its lack of post-translational modifications. "-baculovirus" indicates the extract from Sf9 cells not infected with baculovirus. **b** The NADPH-menadione oxidoreductase activity of wild-type (colored in blue) and the Arg357Cys mutant (colored in gray) *h*MTHFR was carried out at room temperature under varying concentrations of NADPH. Each reaction mixture contained 20 nM of *h*MTHFR. **c** FAD dissociation after dilution of wild-type and mutant *h*MTHFR. Released FAD was detected by fluorometry. The enzyme solution was incubated at 46 °C. The initial rate of FAD release is plotted against enzyme concentration after dilution for the wild-type and mutant enzymes: wild-type (blue), Ala222Val (red), and Arg357Cys (gray). The inset represents the change in fluorescence over time for wild-type and mutant enzymes diluted to 100 nM.

The purified Arg357Cys mutant, which had a typical flavin spectrum, was subjected to FAD release measurement from *h*MTHFR, according to a previously reported method with minor modifications[11]. Briefly, a spectrofluorophotometer RF-5300PC (Shimadzu, Kyoto, Japan) equipped with a cell temperature controller monitored the released FAD from *h*MTHFR. The excitation and emission wavelengths were set at 390 nm and 525 nm, respectively. Concentrated MTHFR was diluted directly into pre-warmed 50 mM KPB at pH 7.2 (3 mL) at 46 ˚C, and the fluorescence intensity of released FAD was monitored for 10 minutes. FAD release from *h*MTHFR$^{R357C}$ was measured across varied concentrations (50~400 nM). At 100 nM, the rate constant of FAD release from the Arg357Cys mutant was 0.030 sec$^{-1}$, which is comparable to that of the wild-type enzyme at 0.031 sec$^{-1}$ (Fig. 3c, inset). The Ala222Val common variant, a well-known thermolabile protein associated with mild hyperhomocysteinemia, was used as a model for the faster FAD release mutant[11]. Consequently, it was evident that *h*MTHFR$^{R357C}$ binds the FAD cofactor as tightly as the wild-type enzyme, while impairing the catalytic function. However, the mutation, located in the linker region away from the active site, posed a challenge in attributing the involvement of the Arg357 residue to one of the catalytic residues.

FAD release from *h*MTHFR is affected by AdoMet, as previously reported[11]. The initial rate of FAD release is decreased in the presence of AdoMet; presumably, in the inhibited state, the T-state, *h*MTHFR resists to release FAD. This gives rise to the idea that the Arg357Cys mutation in the linker region alters the protein conformation to the T-state to inhibit the enzyme activity, rather than affecting the catalytic function as one of the catalytic residues in the active site. The protein conformation of *h*MTHFR$^{R357C}$ is related to the T-state even in the absence of AdoMet, which could explain the properties of the mutant. Taken together, we postulated that the *h*MTHFR$^{R357C}$ patient mutant represents a "T-state locked" enzyme.

### 4. FAD binding site in *c*MTHFR, R-state

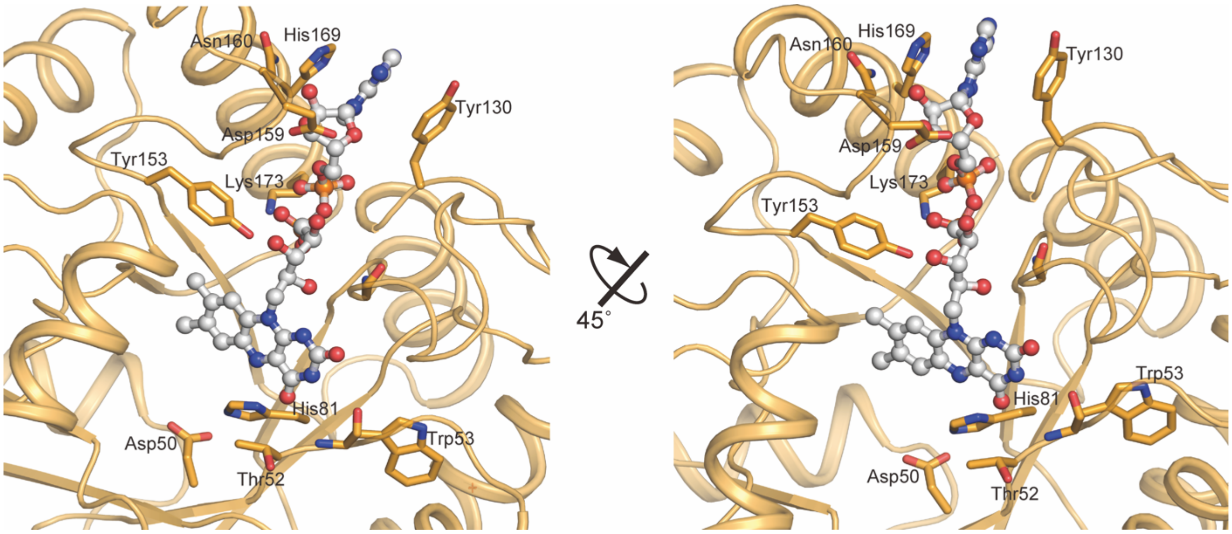


**Supplementary Figure 4. FAD binding site and mode in *c*MTHFR, R-State.** The FAD cofactor of *c*MTHFR is bound in an unoccluded state, making several interactions with the catalytic domain. Of note are the hydrogen bonding networks between N5 of FAD centered on conserved residues His81 and Asp50, along with a strong hydrogen bond between the universally conserved Thr52 and O4 of FAD. His169 and Tyr130 provide π-stacking interactions that serve to hold the adenine moiety in place.

**5. Limited proteolysis of *c*MTHFR in the presence or absence of AdoMet**

Conformational changes of *c*MTHFR, in the presence or absence of AdoMet, were visualized by limited proteolysis followed by SDS-PAGE (Fig. 5a)[12]. Our analysis suggests that the retractable region in the linker (pink, Fig. 5b) should be exposed to the solvent when AdoMet binds to *c*MTHFR. This configuration allows trypsin to digest the linker (Fig. 5c). In contrast, in the R-state, MTHFR securely folds the retractable region between the catalytic and regulatory domains, preventing trypsin cleavage. In the absence of AdoMet, a fraction of *c*MTHFR evades tryptic digestion due to the conformational equilibrium between the R- and T-state. In the presence of AdoMet, *c*MTHFR exhibits increased susceptibility to a lower amount of trypsin, indicating that AdoMet induces a conformational shift to the T-state where trypsin successfully digests the retractable region exposed to the solvent.

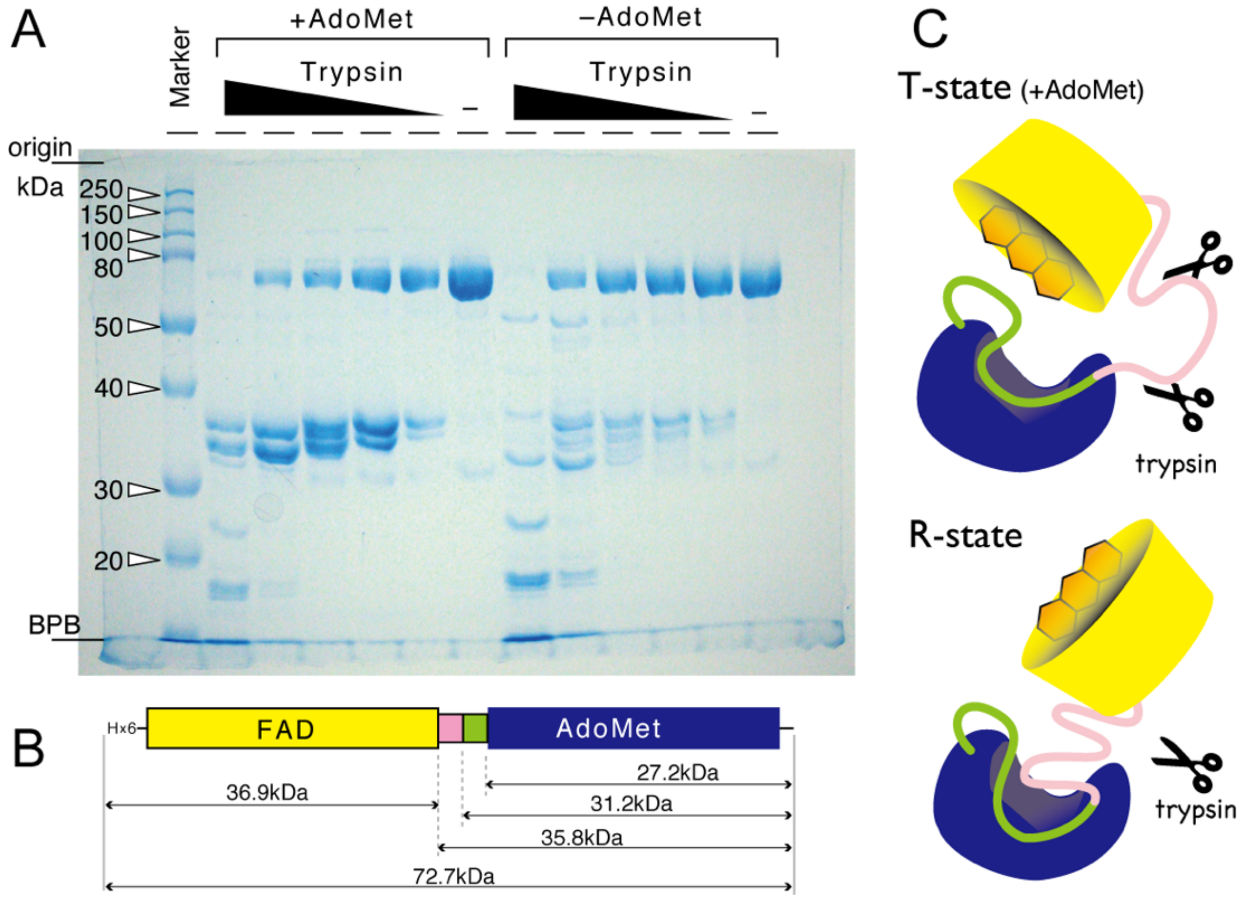


**Supplementary Figure 5. Limited Proteolysis of cMTHFR**. **a** A quantity of 14 µg of purified *c*MTHFR was incubated with varying amounts of trypsin (from ~2 µg to ~0.2 ng) for 20 min at room temperature. The reaction was carried out both in the absence as well as in the presence of 100 µM AdoMet. Trypsin activity was quenched after the reaction by the addition of sample buffer for SDS-PAGE containing 1% SDS followed by heating at 95 ˚C for 10 minutes. The resulting *c*MTHFR fragments were separated by SDS-PAGE and visualized by Coomassie brilliant blue staining. **b** The schematic representation of the domain structure of *c*MTHFR is shown, along with the theoretical molecular weight of each domain. The catalytic domain (yellow) and the regulatory domain (blue) accommodate the FAD cofactor and AdoMet, respectively. Two colors, pink and green, in the linker region represent the "retractable region" and the "inverted cap-for-active-site", respectively. **c** The protein conformations of *c*MTHFR in the T- and R-states are shown in cartoon mode. The color scheme is the same as in panel B. In the R-state, the linker is intricately folded between the catalytic and regulatory domains. Conversely, in the T-state, the retractable region is exposed to the solvent, facilitating trypsin access to the solvent-exposed retractable region, and allowing cleavage of the linker region in the T-state (shown in **c**, top). In contrast, protease access to the linker is difficult when the retractable region is folded in the R-state (shown in **c**-bottom).

### 6. FAD and Tyr361 interaction in T-state

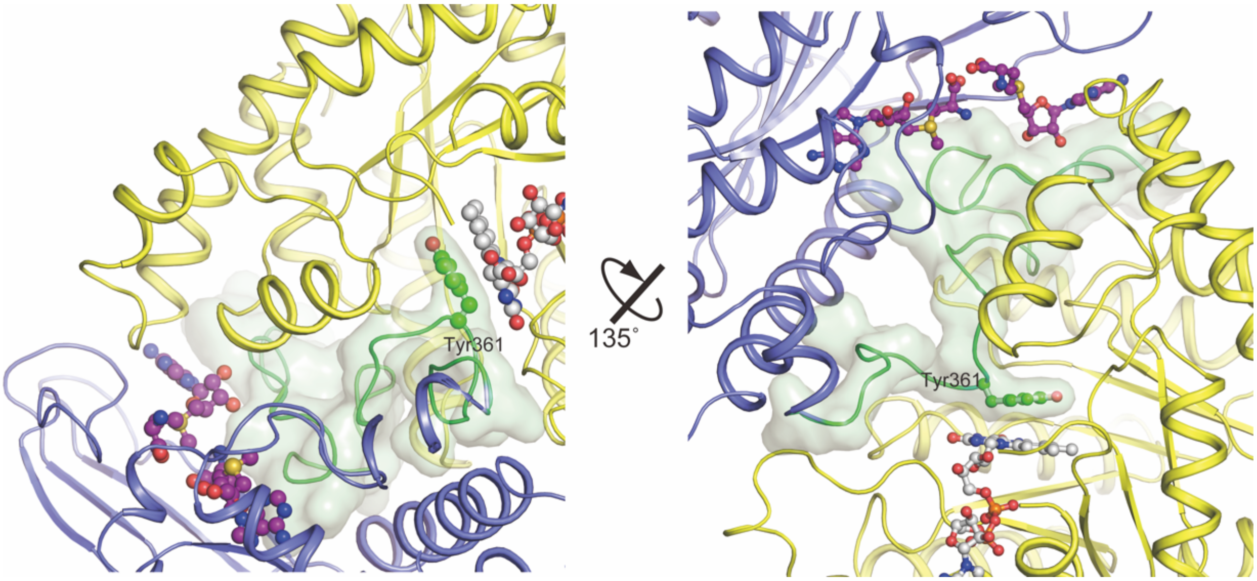


**Supplementary Figure 6. FAD binding site and mode in *c*MTHFR, T-State.** The FAD cofactor of *c*MTHFR is bound in an occluded state. Tyr361 of the velcro-wedge region of the linker provides a π-stacking interaction that serves to hold the FAD isoalloxazine moiety in place and occludes the *si*-face of FAD.

**7. AdoMet binding sites in *c*MTHFR, T-State**

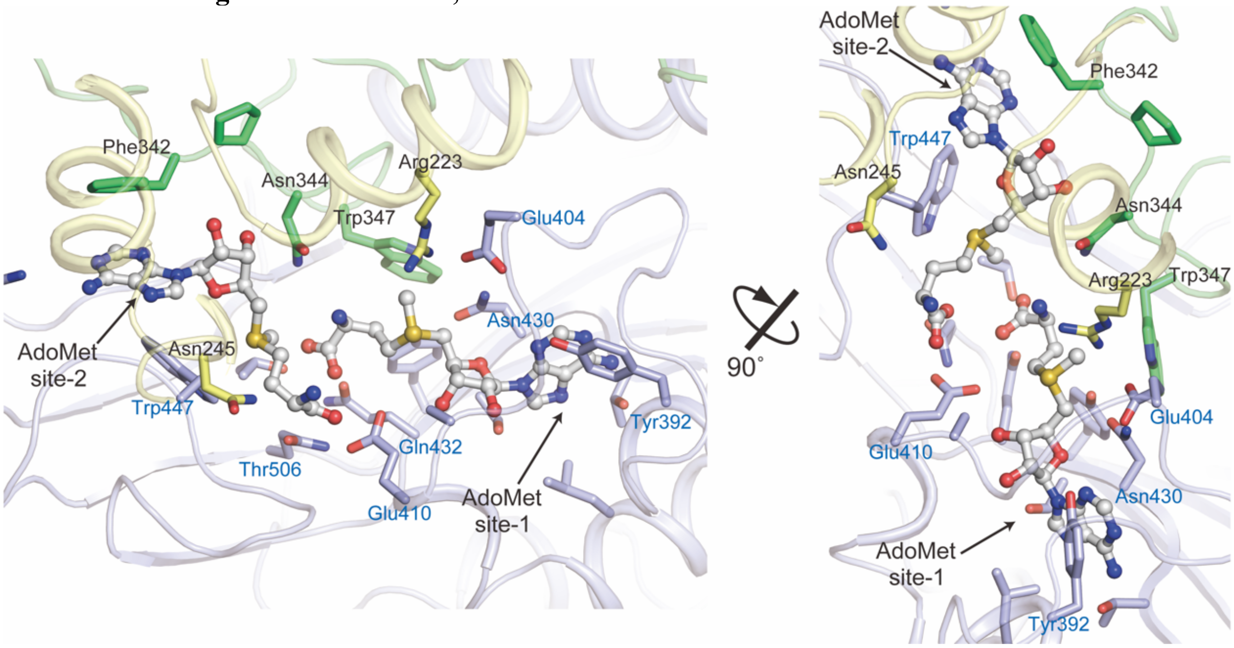


**Supplementary Figure 7. AdoMet binding sites in *c*MTHFR, T-State.** The allosteric inhibitor of *c*MTHFR, AdoMet, is bound in two sites, in the regulatory domain. AdoMet site-1 is the same site AdoHcy occupies in the R-state. AdoMet site-2, the cryptic secondary site, is only unveiled in the T-state, and Phe342 of the velcro-wedge region of the linker and Trp447 of the regulatory domain provide π-stacking interactions that serve to hold the adenine moiety in place.

**8. Electron density around AdoMet allosteric inhibitors in *c*MTHFR, T-State**

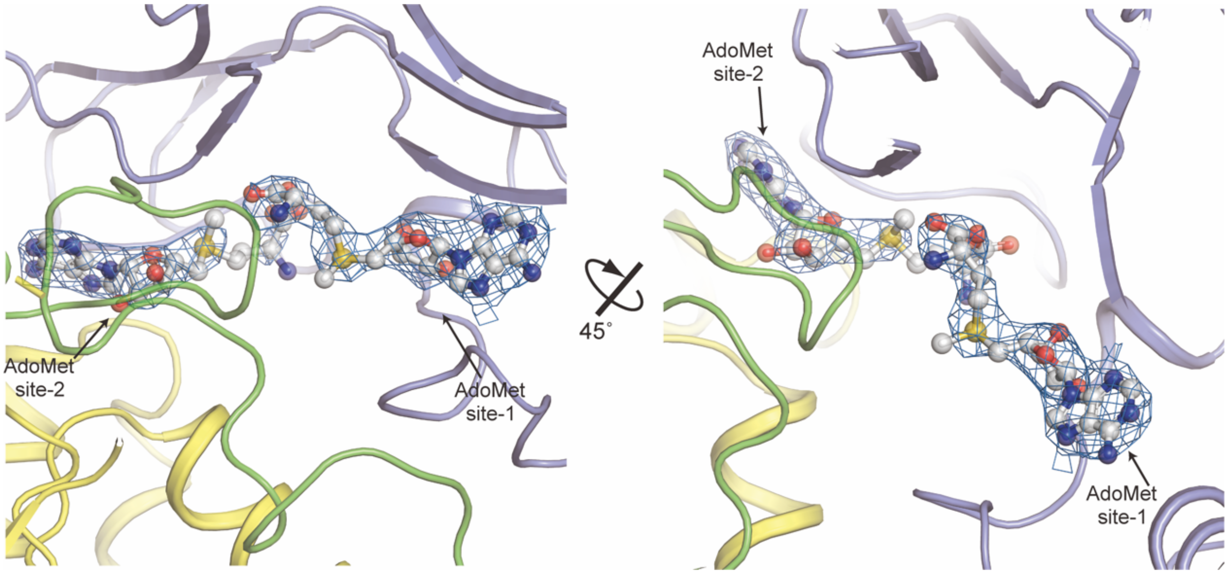


**Supplementary Figure 8. Electron density around AdoMet allosteric inhibitors sites in *c*MTHFR, T-State.** *c*MTHFR, T-State (catalytic domain in yellow and reactivation domain in slate) and the two AdoMet molecules (gray). Their corresponding electron density (2Fo-Fc) contoured at 1.5 s are shown in blue. The allosteric inhibitor of *c*MTHFR, AdoMet, is bound in two sites in the regulatory domain. AdoMet site-1 is the same site AdoHcy occupies in the R-state. AdoMet site-2, the cryptic secondary site, is only unveiled in the T-state.

**9. Structural rigidity of *c*MTHFR domain in R and T states**

Notwithstanding the profound distinctions in protein conformation observed in the R- and T-states of *c*MTHFR structures, the catalytic and regulatory domains exhibited a propensity for rigidity rather than flexibility. The comparative analysis of *c*MTHFR domains in the R-state and the T-state (Fig. 8). The topological configuration of the MTHFR catalytic domain remains conserved across different organisms, ranging from bacterial enzymes to higher eukaryotic enzymes. This domain has the shape of a $\beta_8\alpha_8$ barrel, constituting the TIM barrel fold and comprises approximately 290 amino acid residues. The catalytic domains of *c*MTHFR were aligned in both the R- and T-states (Fig. 9a). The root mean square deviation (RMSD) was calculated to be 0.393, using 221 α-carbon atoms. The structural topology of the regulatory domain presents a unique fold exclusively identified in eukaryotic MTHFR[2], and absent in bacterial MTHFR. The regulatory domains of *c*MTHFR in both R- and T-states were superimposed (Fig. 9b), using 226 residues for the alignment. The RMSD values, based on 203 α-carbon atoms, are impressively low at 0.530. These results underscore the inherent rigidity that characterizes the domain architecture in both the R- and T-states of *c*MTHFR. Consequently, it is suggested that the flexibility inherent in the linker region facilitates the substantial conformational transition observed between the R- and T-state structures.

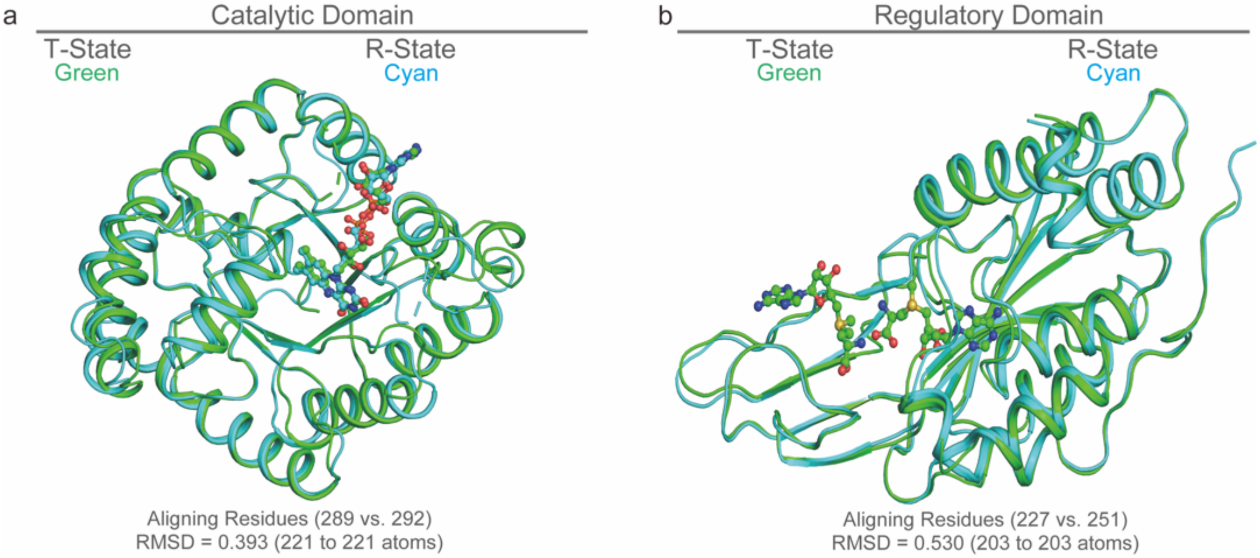


**Supplementary Figure 9. Structural alignment of the catalytic and regulatory domains in the R- and T-states of *c*MTHFR**. **a** The ribbon diagram was used to show the structural alignment of the catalytic domains in *c*MTHFR within the R- and T-states. Cyan and green colors indicate the R and T states, respectively. The FAD cofactors in both structures are shown in stick mode with CMYK colors. **b** The ribbon diagram is used to show the structural alignment of the regulatory domains in *c*MTHFR within the R- and T-states. The color scheme uses cyan for the R-state and green for the T-state. AdoMet ligands bound exclusively in the T-state of *c*MTHFR are shown in stick mode using CMYK colors.

**Supplementary Table 1. X-Ray Data Collection and Refinement Statistics**

| | *c*MTHFR$^{E21Q, L393M, V516F}$ (R-State) | *c*MTHFR$^{R315A}$ (T-State) |
|---|---|---|
| **Data collection** | | |
| Beamline | APS, LS-CAT 21-IDD | APS, LS-CAT 21-IDD |
| Wavelength (Å) | 1.033 | 1.033 |
| Temperature (K) | 100 | 100 |
| Resolution (Å) | 52.04-3.49 (3.62-3.49)* | 75.09-2.83 (2.89-2.83)* |
| Space group | $P2_12_12_1$ | $P22_12_1$ |
| Cell dimensions | | |
| *a*, *b*, *c* (Å) | 117.97, 151.38, 188.05 | 130.66, 149.95, 171.06 |
| α, β, γ (°) | 90, 90, 90 | 90, 90, 90 |
| Observed reflections | 299,056 (32,269) | 555,396 (32,581) |
| Unique reflections | 43,373 (4,481) | 80,491 (4,556) |
| $R_{meas}$ (%) | 16.4 (218.5) | 14.9 (138.3) |
| $R_{merge}$ (%) | 14.3 (191.1) | 12.6 (117.1) |
| $\langle I/\sigma \rangle$ | 8.3 (1.2) | 10.1 (1.9) |
| CC(1/2) | 0.996 (0.562) | 0.987 (0.672) |
| Multiplicity | 6.9 (7.2) | 6.9 (7.2) |
| Completeness (%) | 99.6 (99.9) | 99.8 (100.0) |
| Wilson *B*-factor (Å$^2$) | 50.0 | 52.00 |
| **Refinement** | | |
| Resolution (Å) | 52.04 – 3.49 | 75.09 - 2.83 |
| No. reflections | 41,089 (2,181)‡ | 76,330 (4,071)‡ |
| $R_{work}/R_{free}$ (%) | 26.1/27.2 | 19.3/22.2 |
| No. of non-H atoms | | |
| Protein | 19,081 | 17,836 |
| Water | 4 | 336 |
| Ligand | 212 | 501 |
| B-factors (Å$^2$) | | |
| Protein | 50.0 | 51.3 |
| Water | 80.68 | 61.28 |
| Ligand | 80.68 | 80.98 |
| R.m.s. deviations | | |
| Bond lengths (Å) | 0.0092 | 0.0106 |
| Bond angles (°) | 1.17 | 1.37 |
| Ramachandran Plot | | |
| Favored/allowed/outliers | 97.5/2.4/0.1 | 96.9/2.7/0.4 |
| MolProbity Score | 1.00 ($100^{th}$ percentile) | 1.59 ($100^{th}$ percentile) |
| PDB | 8UY1 | 8UY2 |

* Highest-resolution shell is shown in parentheses.

‡ Number of reflections used for cross-validation

**Supplementary Table 2. Bacterial and insect strains, plasmids, and synthetic oligonucleotides used in this study**

***Strains***

| Cell line | Relevant characteristics | Sources |
|---|---|---|
| *E. coli cells* | | |
| *XL1-Blue* | Routine cloning strain, tetracycline resistance | Stratagene |
| BL21star(DE3) | Widely used T7 expression system, no antibiotic resistance | Invitrogen |
| *Insect cells* | | |
| Sf9 (*Spodoptera frugiperda*) | Host cell for baculovirus-insect cell expression system | Invitrogen |

***Plasmids***

| Names | Relevant characteristics | Ref. or sources |
|---|---|---|
| *Human MTHFR clones* | | |
| pFBHT(hMTHFR$^{wt}$) | wild-type hMTHFR cDNA in pFastBac HT donor vector, encoding an N-terminal His-tag with TEV cleavage site, resistant to ampicillin and gentamicin | Yamada *et al.* 2001[11] |
| pFBHT(hMTHFR$^{A222V}$) | Ala222Val mutant | Yamada *et al.* 2001[11] |
| pFBHT(hMTHFR$^{R357C}$) | Arg357Cys mutant | This Work |
| *Fungal MTHFR clones* | | |
| pMA(cMTHFR$^{wt}$) | synthetic wild-type cMTHFR cDNA in pMA vector, containing the *C. thermophilum* MTHFR cDNA, resistant to ampicillin | GeneArt/Invitrogen (This Work) |
| pMCSG7(cMTHFR$^{wt}$) | wild-type cMTHFR in pMCSG7 vector encoding an N-terminal His-tag with TEV cleavage site, resistant to ampicillin | This Work |
| pMCSG7(cMTHFR$^{R315C}$) | Arg315Cys mutant | This Work |
| pMCSG7(cMTHFR$^{R315A}$) | Arg315Ala mutant | This Work |
| pMCSG7(cMTHFR$^{E21Q, L393M, V516F}$) | Glu21Glu, Leu393Met, and Val516Phe triple-mutant | This Work |

***Oligonucleotides***

| Names | Sequences | Descriptions |
|---|---|---|
| hMTHFR_R357C-f | 5' - CAGTGCACACCCCAAGTGCCGAGAGGAAG - 3' | Site-directed mutagenesis |
| hMTHFR_R357C-r | 5' - CTTCCTCTCGGCACTTGGGGTGTGCACTG - 3' | Site-directed mutagenesis |
| cMTHFR_LIC-f | 5' - TACTTCCAATCCAATGCTATGCATATCCGAGACATGC - 3' | LIC, cMTHFR$^{wt}$ |
| cMTHFR_LIC-r | 5' - TTATCCACTTCCAATGTTAAACTGAGGTCTCCGAAGC - 3' | LIC, cMTHFR$^{wt}$ |
| cMTHFR_E21Q-f | 5' - GCCGTCCTTCTCGTTTCAATACTTCCCGCCCAAGAC - 3' | Site-directed mutagenesis |
| cMTHFR_E21Q-r | 5' - GTCTTGGGCGGGAAGTATTGAAACGAGAAGGACGGC - 3' | Site-directed mutagenesis |
| cMTHFR_R315A-f | 5' - GTCTCTGGGCTTCGGTGCTCGCGGGGAGGATGTCC - 3' | Site-directed mutagenesis |
| cMTHFR_R315A-r | 5' - GGACATCCTCCCCGCGAGCACCGAAGCCCAGAGAC - 3' | Site-directed mutagenesis |
| cMTHFR_R315C-f | 5' - GTCTCTGGGCTTCGGTTGTCGCGGGGAGGATGTCC - 3' | Site-directed mutagenesis |
| cMTHFR_R315C-r | 5' - GGACATCCTCCCCGCGACAACCGAAGCCCAGAGAC - 3' | Site-directed mutagenesis |
| cMTHFR_L393M-f | 5' - CCTCTTCATCCGGTACATGAGAAAGGAAATTGACTAC - 3' | Site-directed mutagenesis |
| cMTHFR_L393M-r | 5' - GTAGTCAATTTCCTTTCTCATGTACCGGATGAAGAGG - 3' | Site-directed mutagenesis |
| cMTHFR_V516F-f | 5' - CCCCGGAAAGGAGATCTTCCAGCCTACCATTGTTGAG - 3' | Site-directed mutagenesis |
| cMTHFR_V516F-r | 5' - CTCAACAATGGTAGGCTGGAAGATCTCCTTTCCGGGG - 3' | Site-directed mutagenesis |
